# Cell Geometry and Membrane Protein Crowding Constrain *Escherichia coli* Growth Rate, Overflow Metabolism, Respiration, and Maintenance Energy

**DOI:** 10.1101/2024.08.21.609071

**Authors:** Ross P. Carlson, Tomáš Gedeon, Mauricio Garcia Benitez, Campbell Putnam, William R. Harcombe, Radhakrishnan Mahadevan, Ashley E. Beck

**Author notes:** Corresponding author: R.P.C.

## Abstract

The rules of prokaryotic cell design remain elusive. Here, a theory is presented for interpreting growth rate, overflow metabolism, respiration efficiency, and maintenance energy flux based on cell dimensions, membrane protein crowding, and metabolism. The theory employs biophysical properties and systems analysis to successfully interpret phenotypes of *Escherichia coli* K-12 strains MG1655 and NCM3722. These strains are genetically similar but differ in surface area-to-volume (SA:V) ratios (∼30%), growth rate on glucose (∼40%), and overflow-inducing growth rates (∼80%). Six predictions were tested and validated using experimental phenomics, proteomics, and mutant data. Analyses did not require assumptions regarding cytosolic macromolecular crowding highlighting the distinct properties of the theory. Cell geometry and membrane protein crowding are significant biophysical constraints of cell biology.

## INTRODUCTION

Bacterial cell geometry, including cell shape and size, is highly regulated and constrains the surface area available for acquiring nutrients and the volume available for synthesizing proteins (1–4). Cell geometry also impacts molecular crowding by fixing the two- and three-dimensional spaces available for macromolecules (5–9). Three-dimensional cytosolic molecular crowding has been analyzed both explicitly and implicitly using computational biology to predict catabolite repression and phenotype shifts including overflow metabolism, the secretion of fermentation products in the presence of electron acceptors (5–7, 10, 11). A quantitative and predictive understanding of the intersection of cellular geometry, two-dimensional membrane protein crowding, and phenotype, however, is not well studied (12–17).

Cell membranes provide a barrier against environmental stressors, retain macromolecules, provide a platform for the selective transport of molecules, and play a crucial role in energy metabolism including respiration (18, 19) (Fig. 1A). The lipid bilayer is a major building block of the membrane. It has a finite capacity to host embedded and adsorbed proteins due to a combination of steric protein crowding effects and the loss of membrane integrity at high protein loading (18, 20–23).

**Figure 1.**
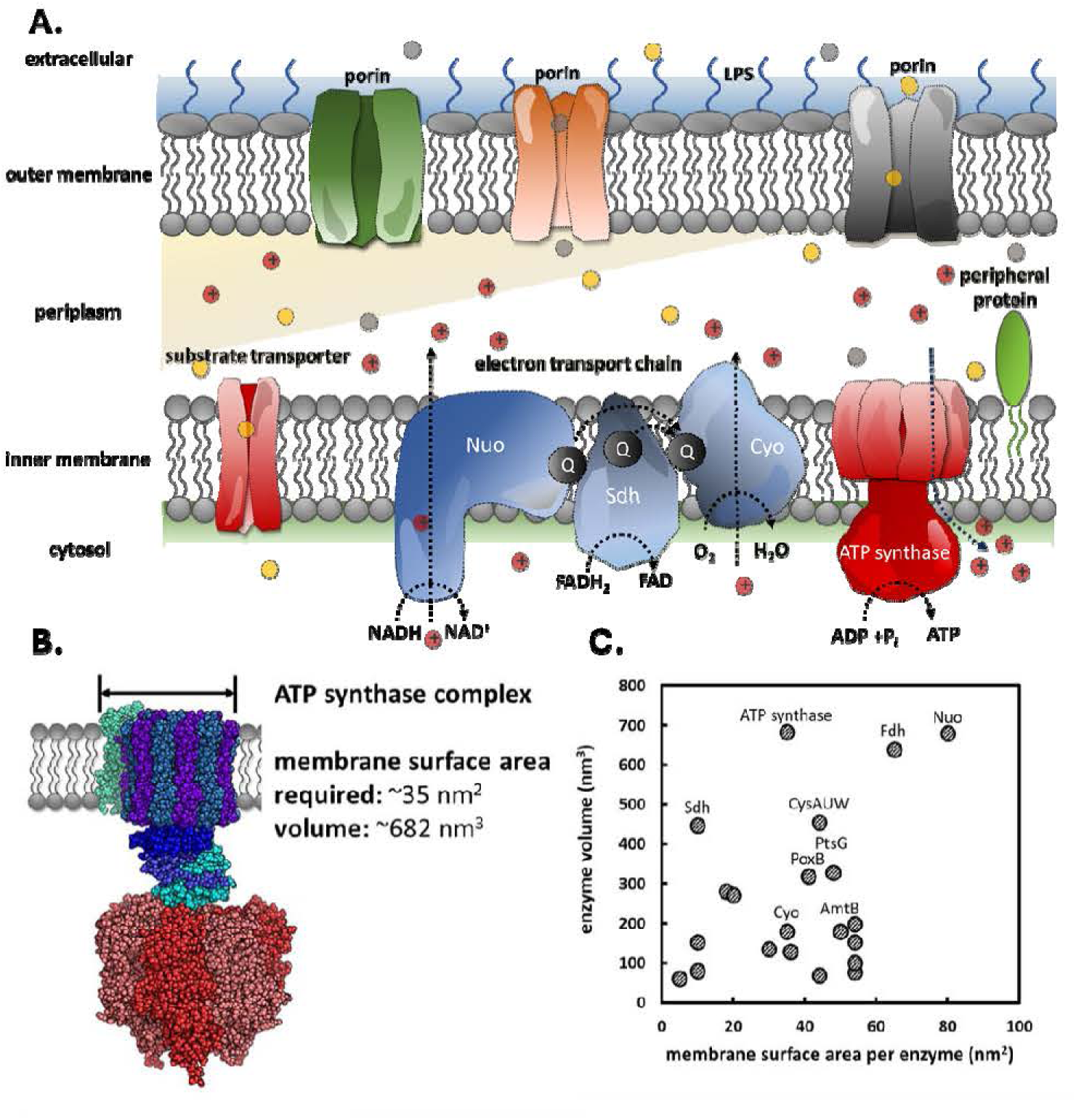
*E. coli* membrane and enzyme properties. **A)** The inner membrane enables critical cellular functions including substrate transport and oxidative phosphorylation. Membrane surface area is finite. **B)** The physical dimensions of membrane-associated enzymes influence both two- and three-dimensional molecular crowding. ATP synthase image modified from Wiki Commons. **C)** Membrane surface area requirements and enzyme volume for 22 central metabolism enzymes do not correlate well. Data available in supplementary data S1 and S3.

Cellular surface area to volume ratios (SA:V) and membrane macromolecular crowding are two foci of the presented work. Systems biology studies have applied enzyme metrics, such as total protein mass, to approximate crowding on a two-dimensional membrane surface, but these approaches generally miss the significant difference between a two- and three-dimensional crowding constraint (10, 19, 24, 25) (Fig. 1B, 1C). The current study makes a distinction between two- and three-dimensional protein crowding on the two-dimensional membrane surface by accounting for biophysical properties of embedded and associated enzymes (10, 24). Enzyme requirements for surface area are not readily predicted from the enzyme mass or volume (Fig. 1B, 1C, supplementary data S1), highlighting the need to study surface properties separately from intracellular crowding.

*Escherichia coli* is a facultative anaerobe and a convenient host for studying tradeoffs between cell geometry, membrane protein crowding, and phenotype. Two well studied and genetically similar *E. coli* K-12 strains, MG1655 and NCM3722, have distinct phenotypes (Fig. 2, supplementary data S1, S2). These cells display exquisite control over their geometry including cell length and width, and we propose that this contributes to the phenotypic differences. The SA:V ratios influence the balance of nutrient and energy fluxes between surface area-and volume-associated processes (1–3). While theoretical studies have addressed the role of surface area on metabolism including Zhuang et al. 2011 and Szenk et al. 2017 (15, 16), no study has developed a quantitative and predictive molecular level theory that accounts for strain-specific differences in SA:V ratios, growth rate dependent changes in SA:V ratios, and growth rate dependent changes in membrane protein crowding (26). Additionally, maintenance energy, which can account for a significant fraction (30 to nearly 100%) of substrate fluxes (27–30), have not previously considered the impact of biophysical constraints like finite membrane surface area and membrane protein crowding. The presented systems biology study demonstrates the remarkable theoretical impact of cell geometry and membrane protein crowding on foundational cellular processes including maximum growth rate, onset of overflow metabolism, respiration efficiency, and ultimately the cellular capacity for energy conservation via maintenance energy generation. These predictions are all accomplished without consideration of cytosolic macromolecular crowding or absolute cellular volume, highlighting the incomplete nature of current cell biology theory. We propose that membrane-associated constraints are complementary to previously described cytosolic-based constraints.

**Figure 2.**
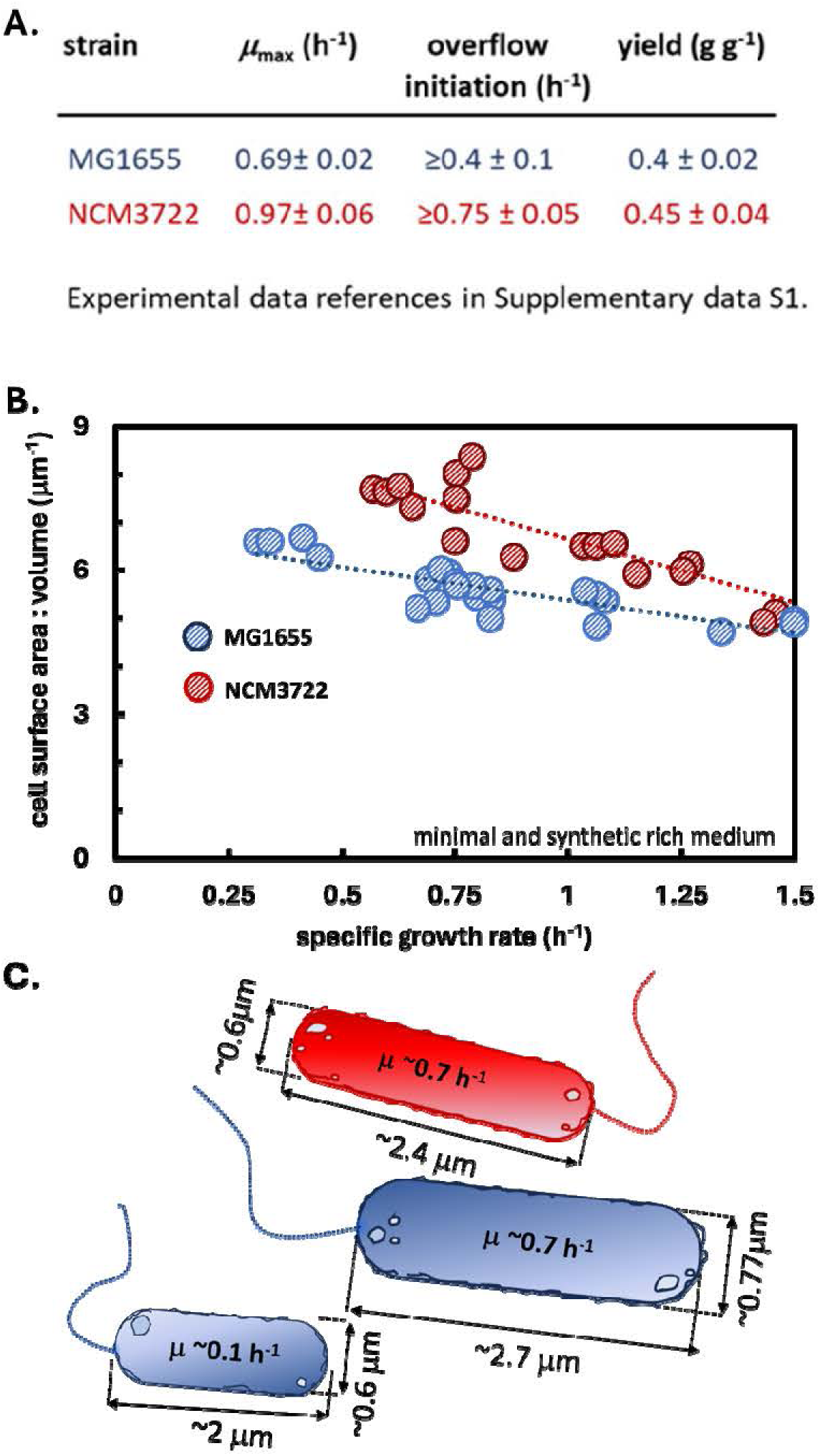
*E. coli* K-12 strains MG1655 and NCM3722 have distinct phenotypes and geometries. **A)** Comparison of strains MG1655 and NCM3722 based on maximum specific growth rate on glucose minimal medium, growth rate at which acetate overflow initiates on glucose minimal medium, and biomass yield on glucose (g biomass (g glucose)^-1^). **B)** Experimental surface area to volume (SA:V) ratios for *E. coli* K-12 strains MG1655 and NCM3722 as a function of specific growth rate. *E. coli* geometry data from Si *et al.* 2017 (2). **C)** Graphical representation of differences in cell geometry (cell diameter and length) as a function of *E. coli* strain (blue = MG1655, red = NCM3722) and specific growth rate. Data available in supplementary data S2.

## MATERIALS AND METHODS

### Enzyme parameters

Computational parameters including enzyme *k*_cat_ (*i.e*. turnover) numbers, enzyme surface area, cellular water content, and cell density for *E. coli* were estimated using literature reviews. Parameter values and references are provided in supplementary data S3. *k*_cat_ numbers were temperature corrected to 37°C using a Q_10_ number of 2 (31). Analyses assumed all membrane-associated enzymes were saturated (ε_i_ = 1) during batch growth, as typical concentrations of medium components during batch growth are ∼2 orders of magnitude greater than the average K_M_ values of *E. coli* enzymes, ∼0.1 mM. During simulation of MG1655 chemostat growth, the saturation parameter (ε_i_) for glucose transport (PtsG) was adjusted based on dilution rate.

### Metabolic model

The *E. coli* metabolic model was based on published models (32, 33). Every model reaction was balanced for atoms and electrons. The biomass reaction was constructed using theory developed by Neidhardt *et al.,* as described previously (32, 34). The biomass macromolecular composition on a dry mass basis was 78% protein, 10% RNA, 6% DNA, and 6% lipid and polysaccharide (34), the elemental composition was C_1_H_1.96_O_0.52_N_0.28_P_0.03_, and the degree of reduction was 4.24 oxidizable electrons per carbon mole of biomass (NH_3_ basis). The biomass ATP demand for monomer synthesis and macromolecule polymerization was 36.9 mol ATP per kg dry biomass (32). The biomass reaction did not account for any maintenance energy requirements as those fluxes were calculated separately. The maximum potential for maintenance energy fluxes (*q_ATP_*) was calculated from experimental data by constraining exchange fluxes to consensus data and maximizing flux through the ATP hydrolysis reaction (reaction identifier: *ATPm*). Thus, the calculated maintenance energy fluxes account for both growth-and nongrowth-associated ATP (GAM, NGAM). The same *E. coli* model was used for both *E. coli* strain MG1655 and NCM3722 simulations while applying strain-appropriate SA:V ratios and consensus data (supplementary data S1, S2, S3, S4). Unless otherwise noted, enzyme saturation (*ε*_i_), enzyme over/under capacity factor (*ω_i_*), and allosteric regulation based on membrane properties (*σ_i_*) were set to 1.

The metabolic model is provided in supplementary data S4. All flux balance analysis simulations used COBRA Toolbox (35) with Gurobi as the solver. FBA is a linear algebra based method of interrogating stoichiometric metabolic models (36). By assuming a (quasi) steady state, FBA identifies a network flux distribution that optimizes an objective function while applying user defined constraints such as bounds on the flux permitted through particular reactions (36). The FBA linear optimization problem is formalized as:

maximize

subject to

where *S* is the stoichiometric matrix and defines the stoichiometry of all metabolites in all reactions, *v* is the vector of reaction fluxes, and *c* is the vector specifying the objective function *Z* such as the flux toward maintenance energy. Fluxes through each reaction *v_i_* were constrained by assigned lower (*lb_i_*) and upper flux (*ub_i_*) bounds (36).

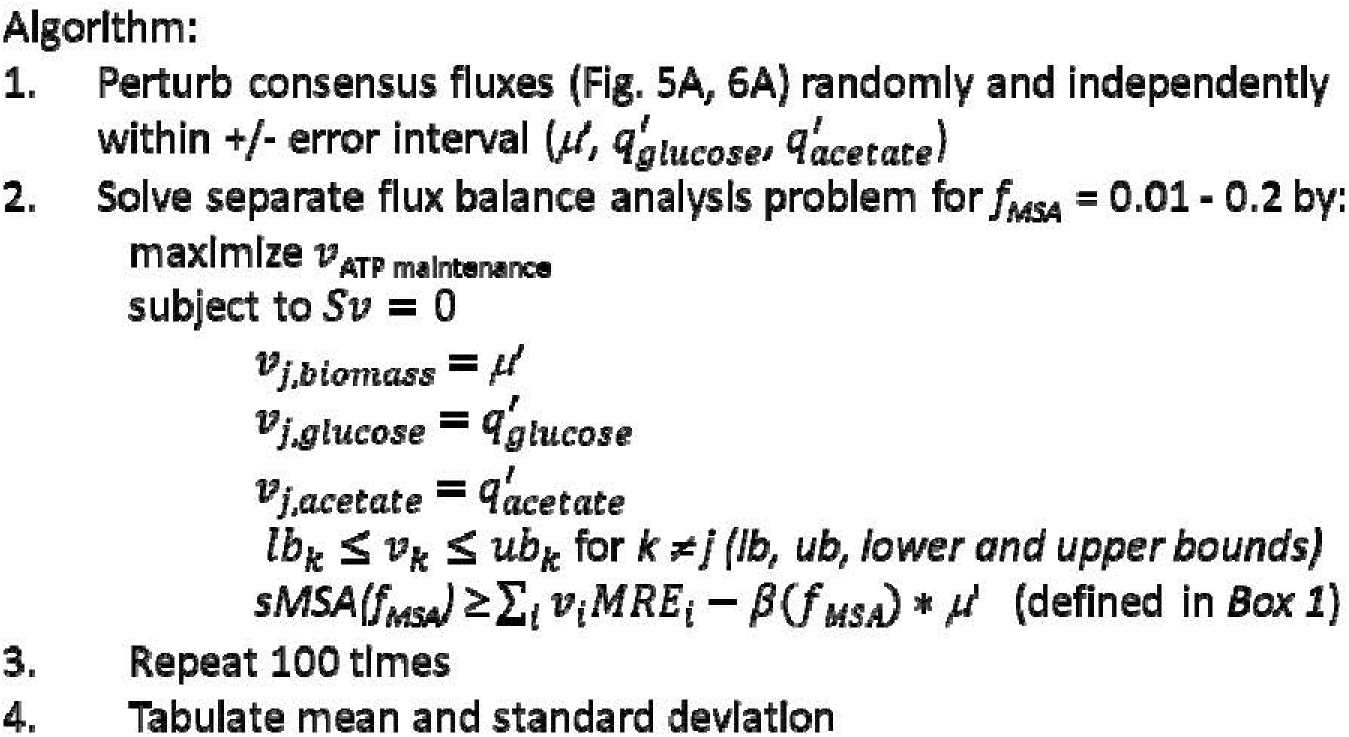

A summary of the code used in the current study can be found in supplementary data S5.

## RESULTS

### *E. coli* K-12 strains MG1655 and NCM3722 have different SA:V ratios and phenotypes

We hypothesize that the biophysical properties of the bacterial inner membrane constrain phenotype and developed a systems biology methodology to test this hypothesis. The methodology was developed using *E. coli* strain MG1655 data, and the predictions were then tested using independent data from *E. coli* strain NCM3722. These two strains are genetically similar but have distinct phenotypes including different maximum growth rates on minimal salts medium, different acetate overflow metabolisms, and different surface area to volume (SA:V) ratios (Fig. 2A, 2B, 2C). Bacterial cells follow “growth laws” and modulate their cell size and growth rate as a function of environment (37, 38). Si et al. measured *E. coli* wild type MG1655 and NCM3722 cellular dimensions and growth rates as a function of different nutrient conditions (2). Cell size measurements are foundational but are technically challenging and can differ substantially between research groups (2, 39, 40). Having a data set of cell dimensions for both strains collected from a single research group builds confidence in the comparisons. SA:V ratios decreased with increasing growth rate because the cell length and width increased (Fig. 2C) (1, 2). NCM3722 cells are smaller than MG1655 at low growth rates, leading to higher SA:V ratios (Fig. 2B, 2C). The different SA:V ratios are proposed to play a role in strain phenotypes. NCM3722 has a specific growth rate on glucose salts media that is ∼40% faster than MG1655 (0.97 ± 0.06 h^-1^ vs. 0.69 ± 0.02 h^-1^, respectively; Fig 2A, supplementary data S1, S2). MG1655 displays an acetate overflow metabolism at growth rates ≥ 0.4 ± 0.1 h^-1^ while NCM3722 displays acetate overflow at growth rates ≥ 0.75 ± 0.05 h^-1^, which are faster than μ_max_ for MG1655 (Fig. 2A, supplementary data S1). Additionally, MG1655 has a cellular volume that is approximately two times larger than NCM3722 at a growth rate of 0.65 h^-1^, whereas MG1655 has a SA:V ratio ∼30% smaller than NCM3722 at this growth rate (Fig. 2B, supplementary data S2). We present evidence that the cell SA:V, as well as the membrane capacity to host proteins, correlate with the phenotypic properties of the strains, suggesting these biophysical properties are important for analyses of cell biology.

### Membrane proteome is dynamic with respect to growth rate and environment

The *E. coli* membrane proteome is dynamic as a function of growth rate and environment (41–45) (Fig. 3A, 3B, 3C). The membrane hosting of central metabolism proteins (protein crowding) including glucose transport, electron transport capacity, and ATP synthase per cell volume increased with growth rate (Fig. 3A) (42–45) (supplementary data S6). In fact, the protein hosting increased faster than the decrease in SA:V (Fig. 2B), resulting in a net gain of membrane catalytic potential per cell volume (protein crowding multiplied by SA:V) with growth rate. Note both sets of data are plotted on a volume basis to facilitate comparisons. The areal densities of central metabolism proteins for *E. coli* K-12 were determined using three proteomics data sets (42–45) and cell geometry (2). Copy number of membrane-associated proteins was converted into an occupied surface area per cell volume using enzyme properties and cell geometry (2, 42–45) (supplementary data S2, S6). The presented trends are for glucose minimal salts media. Faster growth rates were reported in media containing multiple substrates, but the studies did not include detailed substrate uptake data, precluding its inclusion in the current analyses. The presented analyses pooled data for *E. coli* K-12 strains MG1655 and BW25113 which are closely related and have similar growth rates, cell geometries, and phenotypes (39). For simplicity they are collectively referred to here as MG1655.

**Figure 3.**
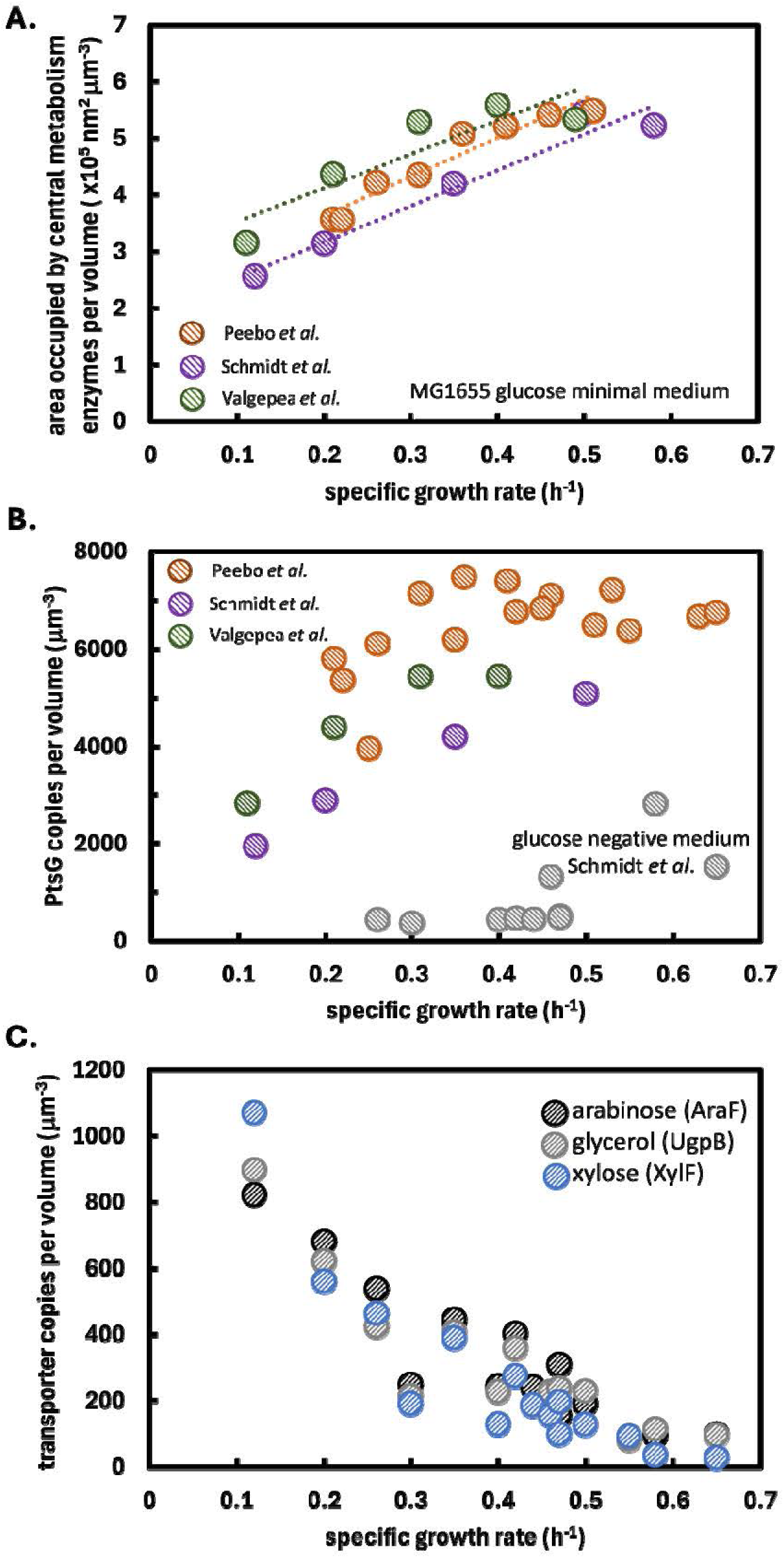
Dynamic membrane proteome of *E. coli* MG1655 as a function of growth rate and environment. **A)** The capacity of the *E. coli* MG1655 inner membrane to host proteins, expressed here as nm^2^ occupied by central metabolism enzymes per volume (μ^-3^). Data from *E. coli* cultures grown on glucose salts medium. Proteomics data from (42–45). Enzyme properties and calculations can be found in supplementary data S1, S3, and S4. Analysis was limited to cultures grown on glucose minimal medium. Calculations were not possible for complex medium because the consumption rates of the numerous substrates were unknown. **B)** Glucose transporter (PtsG) copy number per cell volume as a function of growth rate and growth medium. Gray data points are from glucose negative medium. Proteomics data from (42–45). Supplementary data S6. **C)** Alternative, high affinity sugar transporter copy number per cell volume as a function of growth rate during chemostat growth on glucose containing medium. The specific substrate for each transporter was not present in the medium. Proteomics data from (42–45). Supplementary data S6.

*E. coli* MG1655 synthesized more glucose transporter complexes (PtsG) per volume with increasing growth rate (Fig. 3B). However, *E. coli* cultures grown in the absence of glucose expressed much lower numbers of PtsG complexes. The proteomics data are from chemostat cultures suggesting the cells have a finite surface area and prioritize environmentally relevant membrane proteomes at high growth rates (Fig. 3B). Meanwhile, *E. coli* grown on glucose at low dilution rates synthesized increasing levels of alternative, high affinity transporters for substrate such as arabinose, glycerol, and xylose (Fig. 3C). These substrates were not present in the media and the phenotypes likely represented a hedge strategy preparing the energy limited cells for alternative substrates, should they become available (41, 42) (supplementary data S6). These unused proteins were only synthesized at low dilution rates where we propose there was excess capacity for the membrane to host proteins.

We propose that finite SA:V and membrane capacity to host proteins preclude the simultaneous and constitutive expression of all membrane proteins at all growth rates. The dynamic nature of the membrane proteome is hypothesized to reflect the substantial constraint it places on *E. coli* phenotypes. The remainder of the study presents theory and data to support this proposal.

### Systems biology of cellular SA:V ratios and membrane protein crowding

The foci of this study are the cellular SA:V, inner membrane protein hosting capacity, the areal requirements of inner membrane associated proteins, and the collective role of these biophysical parameters on phenotype. The specific Membrane Surface Area (*sMSA*, units: nm^2^ (g cdw)^-1^) required for membrane-associated enzyme fluxes was quantified using enzyme properties including flux per g cell dry weight (cdw), surface area requirements (nm^2^ enzyme^-1^), and *k*_cat_ (molecules (s enzyme)^-1^). Total *sMSA* required for a phenotype was tabulated by summing the contribution of all active membrane-associated enzymes and by accounting for the growth rate dependent SA:V ratio. Mathematical development is presented in Box 1 while parameters are reviewed in supplementary data S1, S2, and S3. Concisely, the catalytic potential of the cell membrane, on a per g cdw basis, strongly influences what phenotypes are possible. The catalytic potential of the membrane is positively correlated with the cellular SA:V and the membrane protein crowding.

The *sMSA* constraint was evaluated using a metabolic model of *E. coli* (32, 33) and flux balance analysis (FBA) (35). The *sMSA* constraint was implemented by incorporating a membrane surface area requirement for each membrane-associated reaction, termed *MRE_i_* (membrane real estate) (Box 1) (15). Flux through membrane-associated enzymes was constrained by a finite *sMSA* pool which changed in magnitude based on growth rate dependent SA:V ratios and protein crowding (quantified as the fraction of the Membrane Surface Area occupied by central metabolism proteins, *f_MSA_*) (supplementary data S4, Box 1). The *sMSA* balance was enforced concurrently but orthogonally to standard FBA mass balances (mmol (g cdw h)^-1^). The *in silico* theory was termed specific Membrane Surface Area-constrained Flux Balance Analysis (*sMSAc-FBA*) (35). This novel formulation contrasts with previous studies where the weights on the membrane-associated fluxes were the molecular masses of the protein, which correlates with enzyme volume (10, 19, 24, 25, 46), as opposed to the surface area occupied as in the presented work (Fig. 1C). The presented analyses did not consider cytosolic protein investment nor cytosolic macromolecular crowding to isolate the influence of the membrane catalytic constraint on phenotypic predictions.

### *E. coli* MG1655 growth rate and biomass yield are constrained by membrane surface area, protein crowding, and maintenance energy

Cell growth requires fluxes of nutrients across the membrane for biomass synthesis and for maintenance energy generation. Intracellular reactions for macromolecular synthesis are well characterized while maintenance energy requirements are still largely a black box (47). Maintenance energy is cellular energy consumed for functions other than the direct production and polymerization of biomass components (48). The theoretical impact of finite *sMSA* on these fluxes was quantified for *E. coli* strain MG1655 growing on glucose salts medium (Fig. 4A). Briefly, a matrix of biomass synthesis rates (*q_x_*) and maintenance energy fluxes (*q_ATP_*, aggregate of growth- and nongrowth-associated maintenance energy (GAM, NGAM, respectively)) was defined and applied as FBA flux constraints. Each parameter combination comprised of a unique *q_x_* and *q_ATP_*, while an *sMSA* optimized phenotype was identified by minimizing glucose uptake (*f_MSA_* = 0.07, Fig. 4A, Box 1). This analysis used an ‘unbiased’ approach to maintenance energy fluxes to illustrate the effect of *sMSA* on *in silico* phenotypes. Maintenance energy magnitude is largely a parameter fitting exercise requiring substantial assumptions like respiration efficiency; additionally, the magnitude of maintenance energy changes with environment (29, 49–51). For example, *E. coli* cultures grown aerobically on glucose under phosphate or iron limitation or grown anaerobically on glucose appear to have substantially different maintenance energy fluxes than reference, aerobic glucose-limited *E. coli* cultures (52–55). With the data presented, a reader can apply a maintenance energy vs. growth rate relationship of their choosing to recreate a metabolic solution space of interest. (supplementary data S7). The plot uses an *f_MSA_* of 0.07 because, as shown below, this value corresponds with the *E. coli* K-12 maximum growth rate on glucose minimal medium and thus represents a convenient reference point. The supplementary material presents MATLAB code to recreate the analysis with any *f_MSA_*value from 0.01 to 0.2 (supplementary data S5).

**Figure 4.**
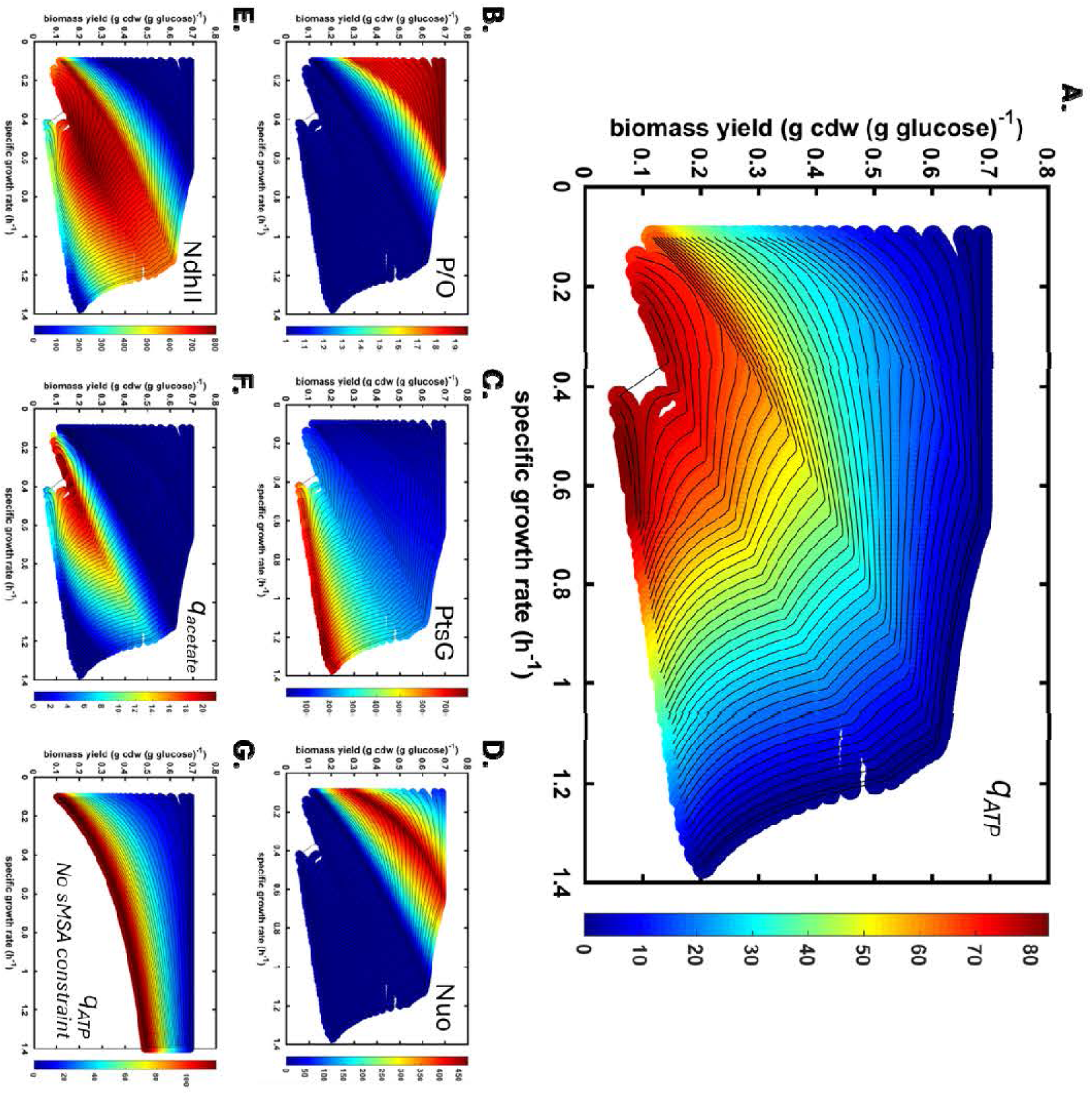
Predictive growth rate – biomass yield phenotype space for *E. coli* strain MG1655 constrained by membrane surface area and membrane protein crowding. **A)** Maintenance energy flux (mmol ATP (g cdw h)^-1^) shown by the color gradient (protein crowding, *f_MSA_* = 0.07). Black contour lines are maintenance energy isoclines. Maintenance energy flux accounts for ATP in excess of ATP used for macromolecule polymerization. **B)** Color gradient represents the P/O number which quantifies the efficiency of oxidative phosphorylation. **C)** Color gradient quantifies the areal density of glucose transporter PtsG (complexes μm^-2^). **D)** Color gradient quantifies areal density of the electron transport chain component Nuo (complexes μm^-2^). **E)** Color gradient quantifies areal density of the electron transport chain component Ndh II (complexes μm^-2^). **F)** Color gradient quantifies overflow metabolism *q*_acetate_ (mmol acetate (g cdw h)^-1^). **G)** Negative control simulation unfettered by the *sMSA* constraint. Color gradient quantifies potential for maintenance energy generation (*q_ATP_*). *f_MSA_* = fraction of membrane surface area.

Metabolic processes competing for a finite pool of *sMSA* resulted in rate-yield tradeoffs (16, 56). Phenotypes with high biomass yields (g biomass (g glucose)^-1^) required larger investments of *sMSA* for high efficiency electron transfer chain (ETC) components, quantified by the P/O number (Fig. 4B). The P/O number is the ratio of ATP produced via respiration per two electrons transferred to O_2_. *E. coli* has the genetic potential to operate ETC configurations with P/O numbers from ∼0 to 2 (57–59). ETC efficiency is not influenced by substrate-level phosphorylation such as ATP synthesis associated with acetate overflow.

Phenotypes with low growth rates and small maintenance energy fluxes can support high P/O numbers because of the modest *sMSA* requirements for transporters and ATP synthase complexes (upper left Fig. 4B, supplementary data S7). Phenotypes with high growth rates require larger investments of *sMSA* for glucose transporters (PtsG) (Fig. 4C, supplementary data S7). These tradeoffs generate gradients in other phenotypic properties including the potential for maintenance energy fluxes (*q*_ATP_) (Fig. 4A), use of parallel ETC NADH dehydrogenase enzymes Nuo (Fig. 4D) or Ndh II (Fig. 4E), and acetate overflow metabolism (*q*_acetate_) (Fig. 4F). Secretion of acetate generates ATP via substrate-level phosphorylation and reduces the need for *sMSA*-intensive ETC components and ATP synthase complexes (15, 16). Presented *q*_ATP_ accounts for the capacity to generate the sum of growth- and nongrowth-associated maintenance energy (GAM and NGAM, respectively) but does not include ATP required to polymerize monomers into macromolecules for biomass synthesis as this energetic cost is included in the biomass synthesis reaction (supplementary data S4).

As a negative control, the same matrix of biomass synthesis fluxes and maintenance energy fluxes was analyzed without the sMSA constraint (Fig. 4G). Without the constraint, there was no rate-yield tradeoff, no maximum growth rate was identified, and the simulations permitted maintenance energy fluxes more than 10-fold higher than with the *sMSA* constraint at fast growth rates.

### μ_MAX_ is a Pareto tradeoff between biomass production and maintenance energy generation

Finite *sMSA* may necessitate phenotypic tradeoffs, providing a new lens with which to interpret maximum growth rate. Consensus extracellular fluxes were assembled for *E. coli* strain MG1655 growing on glucose salts media (Fig. 5A, supplementary data S1). Data span growth rates from 0.1 h^-1^ to *μ_max_* (6, 44, 46, 60, 61). *sMSA* theory was applied integrating the experimental fluxomics data (Fig. 5A), with growth rate dependent changes in: SA:V, protein crowding (*f_MSA,_* 0.02-0.09), and PtsG enzyme saturation (Fig. 5B). These data sets were all applied as constraints to *sMSAc-FBA* analyses which optimized the maximum possible maintenance energy generation (*q_ATP_*) as a function of growth rate (Fig 5B, 5C, 5D). *q_ATP_* was calculated 100 times for each growth rate by randomly and independently perturbing each experimental consensus flux within an assumed error interval (±7% of flux) and using these perturbed fluxes as the *sMSAc-FBA* reaction constraints. Data points in Fig. 5C – 5E show the mean from 100 simulations; error bars represent the standard deviation.

**Figure 5.**
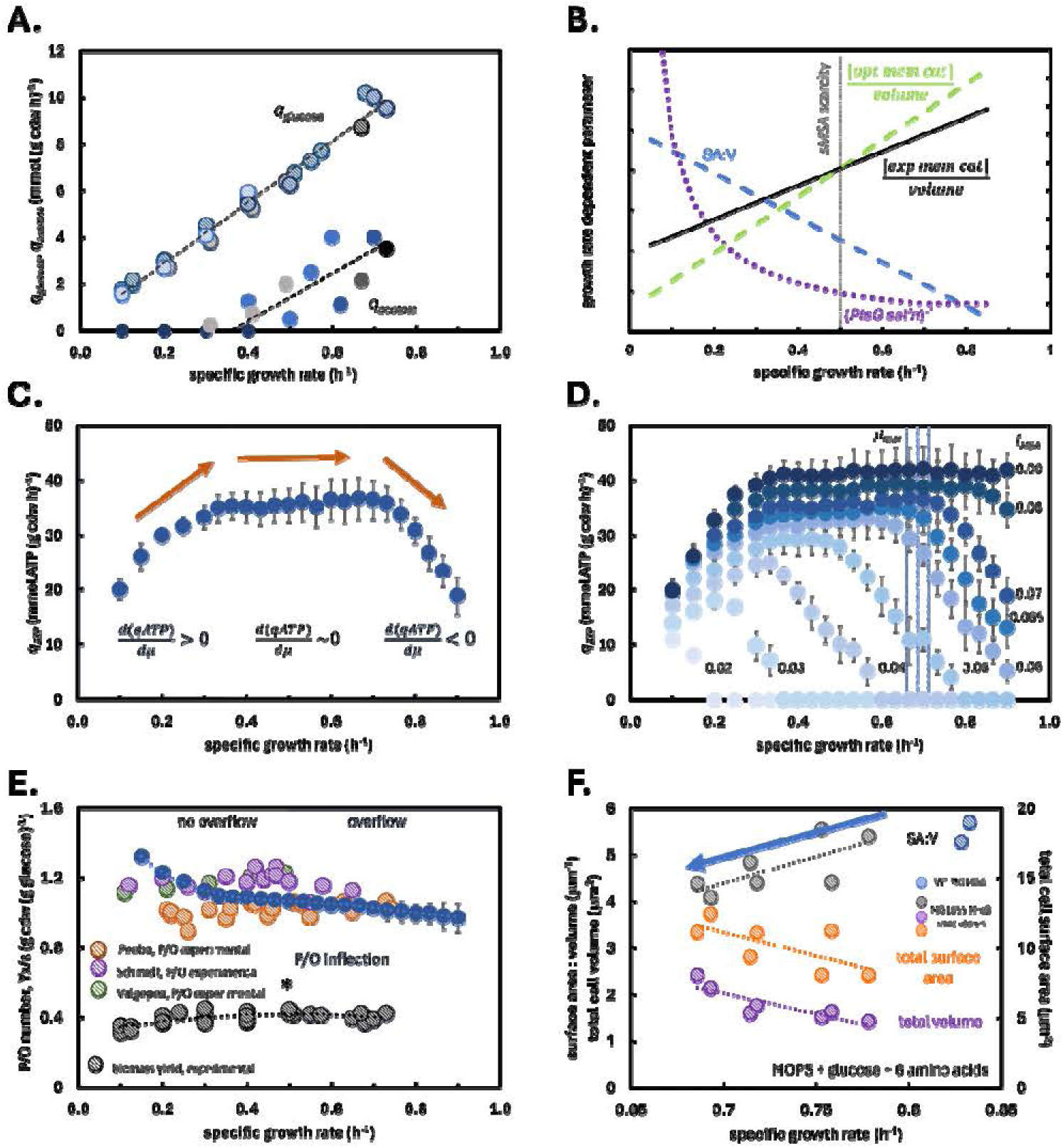
*E. coli* MG1655 phenotypic properties constrained by membrane surface area and protein crowding. **A)** Experimental data from *E. coli* K-12 strain MG1655 used to build consensus fluxes as a function of growth rate. Experimental data references can be found in supplementary material S1. **B)** Summary of biological processes integrated in *sMSA* analysis including growth rate dependent changes in SA:V, PtsG glucose saturation, and membrane catalytic capacity per volume, optimal (opt mem cat) and experimental (exp mem cat). Error bars represent standard deviation of 100 simulations. **C)** An example of the nexus of experimental consensus fluxes and *sMSA* theory defines three regions based on the slope between the capacity to support biomass fluxes and maintenance energy fluxes. The negative slope defines a Pareto tradeoff. Error bars represent standard deviation of 100 simulations. **D)** Maintenance energy potential of the *E. coli* MG1655 consensus fluxes as a function of membrane protein crowding quantified by *f_MSA_*. A series of simulations was performed enforcing different membrane protein crowding (*f*_MSA_ = 0.02-0.09, or 2-9% of occupied membrane area). Error bars represent standard deviation of 100 simulations; see Figure 5B and main text. A Pareto tradeoff between growth rate and maintenance energy flux occurs at the experimental *μ_max_* for an *f_MSA_* of 0.07. Shaded area quantifies experimental maximum growth rate and standard deviation. **E)** Predicted *E. coli* MG1655 P/O numbers (solid blue points) plotted with experimentally determined P/O numbers (striped points) as a function of growth rate. Predicted P/O numbers have an inflection point denoting a change in scarcity of the membrane surface area. Overflow metabolism occurs at severe surface area scarcity, right of inflection point. The P/O number inflection corresponds with higher experimental biomass yields on glucose. Biomass yield maximum highlighted with (*). Error bars represent standard deviation of 100 simulations using strain-specific consensus fluxes. Experimental proteomics data from (42–45). **F)** Experimental testing of *sMSA* theory using an MG1655 strain engineered for titratable MreB expression to control cell SA:V ratios. Decreasing strain SA:V ratio resulted in slower growth rates on glucose medium supplemented with amino acids (gray data). The total volume (purple data) and total surface area (orange data) of the MG1655 strain expressing titratable levels of MreB decreased with growth rate. Data from (2). *f_MSA_* = fraction of membrane surface area. *sMSA* = specific membrane surface area (nm^2^ gcdw^-1^). SA:V = surface area to volume.

Positive slopes between growth rate and maintenance energy flux (Fig. 5C) indicate that both rates can theoretically increase simultaneously without penalty as the membrane can accommodate a balanced increase of substrate transporters, ETC complexes, and ATP synthase complexes. A slope of zero indicates that the growth rate can increase with no reduction in *q*_ATP_; the membrane hosts more substrate transporters while holding the other processes constant. A negative slope indicates that the growth rate can increase only with a concurrent decrease in *q*_ATP_; the increased substrate transporter demand requires a decrease in ETC components and ATP synthase complexes. Negative slopes define a Pareto front between simultaneous increases in growth rate and *q*_ATP_ (Fig. 5C, 5D) (62).

Membrane protein crowding (*f_MSA_*) has the potential to constrain the energetic efficiency of respiration by limiting the ability of the cell to extract useful energy from substrate. For example, there was a ∼4-fold increase in predicted MG1655 maintenance energy potential (10 ± 3 vs. 41 ± 4 mmol ATP (g cdw h)^-1^) for the same consensus fluxes (*μ* = 0.5 h^-1^, *q_glucose_* = 7.2 ± 0.5 mmol (g cdw h)^-1^*, q_acetate_* = 1.7 ± 0.1 mmol (g cdw h)^-1^) when the enforced protein crowding increased from 4% to 9% (*f_MSA_* = 0.04, 0.09) (Fig. 5D, supplementary data S7). The difference in *q_ATP_* potential is based on the availability of *sMSA* required to use higher efficiency ETC components and assemble more ATP synthase complexes. We hypothesize that *E. coli* operates at a maximum membrane protein capacity as this would significantly increase the possible maintenance energy generation, thereby improving fitness.

Experimental *E. coli* MG1655 cultures have a *μ_max_* of 0.69 ± 0.02 h^-1^ on glucose salts media (Fig. 2A, blue shaded area, Fig. 5D) which coincides with the start of a negative slope between growth rate and *q*_ATP_ (*f*_MSA_ = 0.07). The predicted *f_MSA_* = 0.07 is lower than values estimated by Szenk *et al*. (15). The current study adjusted enzyme *k_cat_*values to a temperature of 37°C, resulting in some faster *k_cat_*values and therefore smaller *f_MSA_* requirements (31). All reactions in the *in silico* model were balanced for atoms and electrons (supplementary material S4).

Therefore, all simulations necessarily balanced consensus fluxes with appropriate CO_2_ (necessary for carbon balance) and O_2_ (necessary for electron balance) fluxes as no other electron acceptors were available in the simulations (63).

### Submaximal P/O numbers optimize energy conserving potential of finite membrane surface area

*E. coli* has a branched ETC that can convey electrons using multiple enzymatic routes to acceptors including O_2_ (57–59). These enzymes can conserve varying amounts of substrate chemical energy by converting it into chemiosmotic energy stored as a proton motive force. As mentioned previously, *E. coli* ETC components can operate at P/O numbers between ∼0 and 2 (57–59). P/O number can significantly influence maintenance energy calculation (50). *E. coli* cultures typically operate at submaximal P/O numbers in the range of 1 to 1.5 even at O_2_ sufficiency (reviewed in supplementary data S3). Few theories have been proposed to explain this phenotype (64, 65). We hypothesize *sMSA* theory is relevant to the experimentally observed submaximal P/O numbers.

Data from four proteomics studies (42–45) and enzyme *k*_cat_ numbers were used to calculate experimental P/O numbers as a function of growth rate (Fig. 5E, supplementary data S7, S8). Theoretical growth rate dependent P/O numbers were calculated using *sMSAc-FBA* from experimental consensus fluxes, SA:V ratios, and growth rate dependent membrane protein crowding (Fig. 5E, supplementary data S7, S8). *In silico* P/O numbers at *μ_max_* were comparable to the experimental values (1.04 ± 0.01 and 1.01-1.07, respectively). We propose the experimentally observed submaximal P/O numbers represent an optimized use of the finite *sMSA* that enables optimal substrate fluxes for growth and cellular energy generation and now present additional evidence for this conclusion. Predicted P/O numbers increase rapidly as growth rate decreases. We propose slow-growing cells are not substantially constrained by *sMSA* as the slow growth rate does not require as many glucose transporters, ETC components, or ATP synthase complexes (Fig. 5B). Slow growing cells have sufficient *sMSA* to permit the expression of alternative substrate transporters such as those shown in Fig. 3C.

### Membrane surface area and membrane protein crowding predict onset of overflow metabolism

The intersection of metabolism, membrane protein crowding, and cellular geometry predicts the onset of acetate overflow based on optimization of finite *sMSA* (Fig. 5B). The presented analysis is designed to further test the *sMSA* theory by highlighting its potential significance as a mediator of major phenotypic strategies. *sMSAc-FBA* was used to calculate growth rate dependent P/O numbers for strain MG1655 using the experimental consensus fluxes, SA:V, and growth rate dependent membrane protein crowding (Fig. 5A, Box 1, supplementary data S8).

The theoretical P/O number vs. growth rate curve was fit empirically (analyses of different empirical fittings are presented in supplementary data S8). The inflection point was determined from the second derivative with respect to growth rate. The inflection point identified the growth rate where *sMSA* became sufficiently limiting to warrant the overflow metabolism. The inflection point at μ = 0.5 h^-1^ correlates well with the experimental values for overflow onset, 0.4 ± 0.1 h^-1^ (Fig. 2A, supplementary data S1). At growth rates less than the inflection point, the analysis suggests cells have sufficient *sMSA* to operate substrate transporters, ETC apparatus, and ATP synthase without requiring the overflow tradeoff (Fig. 5B, 5E). The P/O number inflection also coincides with the maximum experimental biomass yields (black circles), providing a basis for defining an optimal biomass phenotype (Fig. 5A, 5E, supplementary data S1).

### Modulating E. coli MG1655 SA:V alters maximum growth rate in predictable manner

Si *et al*. (2) examined the effect of MG1655 SA:V on batch growth rates. A titratable MreB system was used to modulate *E. coli* geometry by changing the cell diameter and cell length and therefore changing SA:V ratios (1–3, 66). The batch growth rate on defined glucose medium supplemented with six amino acids was measured as a function of titrated SA:V. These experiments modulated the cell SA:V using the transcriptome while using the same growth medium (MOPS + glucose + 6 amino acids). In contrast, the data in Figure 2B was collected from wild type cultures subjected to different nutrient environments which provided a different mechanism of controlling growth rate and SA:V (37, 67). Cells with higher SA:V ratio grew faster than cells with lower SA:V ratios, supporting the *sMSA* theory as the theory correlates higher membrane catalytic activity per g cdw (SA:V\**f_MSA_*) with faster substrate uptake, faster growth, and more efficient respiration (Fig. 5F). Interestingly, when the same data set was plotted as total cell surface area or total cell volume vs. growth rate, a negative relationship was observed (Fig. 5F) (1, 3, 68). The same study also analyzed a strain with a titratable FtsZ system. The strain did not achieve a substantial range of SA:V ratios, and the specific growth rates during batch growth were largely unchanged (supplementary data S2)

### Testing sMSA predictions using data from E. coli strain NCM3722

*sMSA* theory, developed using *E. coli* strain MG1655 data, predicted several phenotypic properties based on SA:V ratios and membrane protein crowding (*f_MSA_*). We propose if these phenotypic properties can be predicted for another *E. coli* strain, such as strain NCM3722, then the theory holds meritorious promise. Synthesizing the theoretical framework, we predict the following behaviors will hold for *E. coli* strain NCM3722: **prediction 1,** *sMSAc-FBA* using an *f_MSA_* of 0.07 will predict *μ_max_* at the Pareto front between biomass synthesis and maintenance energy generation; **prediction 2,** *sMSAc-FBA* will only predict an accurate *μ_max_* when biologically relevant SA:V ratios are applied to the analyses; **prediction 3,** *sMSAc-FBA* will quantitatively predict P/O ratios highlighting the optimal use of finite membrane surface area and protein crowding; **prediction 4,** *in silico* P/O ratios, plotted as a function of growth rate, will have an inflection point that corresponds with the experimental onset of acetate overflow; **prediction 5,** the most efficient biomass yield on glucose will correspond with the inflection point of P/O number vs. growth rate, providing a basis for defining an optimal glucose phenotype; and **prediction 6**, culture growth rates will decrease in a fixed nutrient environment when the cellular SA:V ratios are decreased. These evaluations of *sMSA* theory used the same metabolic model with the same enzyme parameters as applied for MG1655 simulations. However, strain-specific SA:V ratios (Fig. 2B) and consensus fluxes were applied (Fig. 6A).

**Figure 6.**
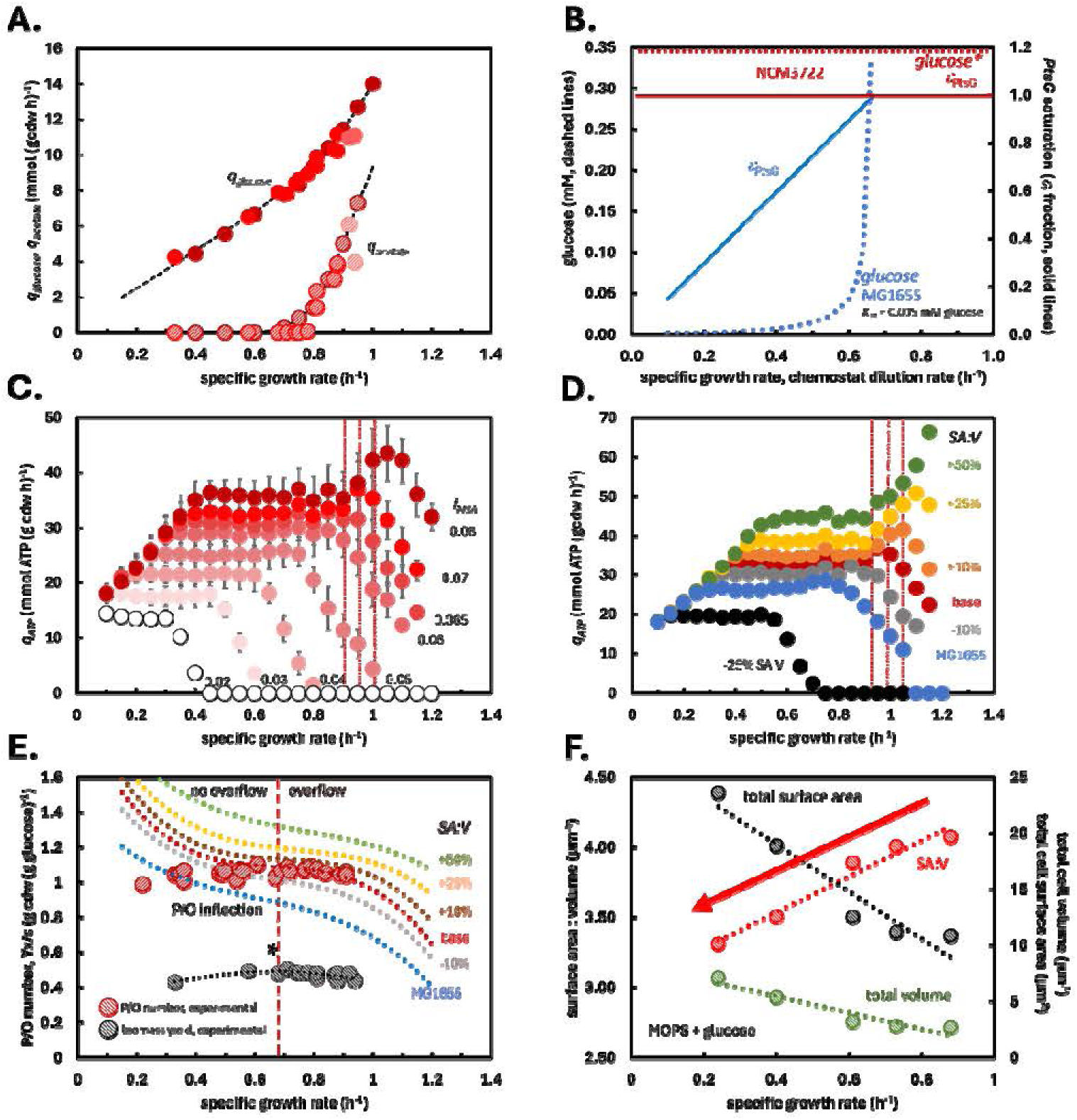
Testing *sMSA* predictions using data from *E. coli* NCM3722. **A**) Experimental data from *E. coli* K-12 NCM3722 used to build consensus fluxes as a function of growth rate. Experimental data references can be found in supplementary material S1. **B**) Relationship between glucose transporter (PtsG) saturation (dotted lines) and as a function of growth rate for wild type MG1655 and recombinant NCM3722. Steady state glucose concentration vs. growth rate for wild type MG1655 and representative batch growth glucose concentration for recombinant NCM3722 cultures. NCM3722 cultures started at approximately 11 mM (off the chart). **C)** Maintenance energy potential of the *E. coli* NCM3722 consensus fluxes as a function of membrane protein crowding quantified by *f_MSA_*. A series of simulations was performed enforcing different membrane protein crowding (*f*_MSA_ = 0.02-0.08, or 2-8% of occupied membrane area). Error bars represent standard deviation of 100 simulations; see Figure 5B and main text. A Pareto tradeoff between growth rate and maintenance energy flux occurs at the experimental *μ_max_* for an *f_MSA_* of 0.07. Shaded area quantifies experimental maximum growth rate and standard deviation. **D)** Effect of applied SA:V ratios on the relationship between biomass synthesis and maintenance energy potential. Simulations fixed *f_MSA_* at 0.07 and varied the SA:V from the experimentally determined relationship (±25% of the experimental values and the MG1655 SA:V relationship). Changing the applied SA:V changed the predicted *μ_max_*. Data points represent the mean of 100 simulations. **E)** Predicted P/O numbers as a function of varying SA:V ratios and growth rate plotted with experimentally determined P/O number. Predicted P/O numbers have an inflection point denoting the change in scarcity of the membrane surface area. Overflow metabolism occurs at severe surface area scarcity, right of inflection point. The P/O number inflection corresponds with higher experimental biomass yields on glucose. Biomass yield maximum highlighted with (*). Experimental proteomics data from (61). **F)** Experimental testing of *sMSA* theory using an NCM3722 derived strain engineered for titratable protein expression which controlled cell volume and SA:V ratios. Decreasing strain SA:V ratio slowed growth rates on glucose medium (red data). The total cell volume (green data) decreased with increasing growth rate. Data from (70). *f_MSA_* = fraction of membrane surface area. *sMSA* = specific membrane surface area (nm^2^ gcdw^-1^). SA:V = surface area to volume.

*E. coli* NCM3722 data were assembled to develop consensus fluxes (6, 46, 69). The majority of the NCM3722 data were derived from a recombinant strain with a recombinant glucose uptake system (6). Briefly, the strain had the native PtsG glucose transport component deleted, and in place, a plasmid-based genetic construct with a titratable PtsG protein was used to control growth rate. The recombinant NCM3722 strain was grown in batch, and the glucose transporter was saturated for all growth rates based on the recombinant design (enzyme K_M_ ∼ 5μm, initial medium glucose concentration S ∼ 11 mM) (Fig. 6B). The recombinant NCM3722 phenotype was applied to the *sMSAc-FBA* simulations using a PtsG enzyme saturation factor _PtsG_ = 1 (Fig. 6B). This was in contrast with data for MG1655 which was collected from wild type cultures with growth rates controlled by substrate concentration modulated by chemostat dilution rates (6, 44, 46, 60, 61). MG1655 simulations accounted for changing PtsG enzyme saturation ( _PtsG_) as a function of growth rate (Fig. 6B).

#### Testing prediction 1

NCM3722 data demonstrated trends analogous to MG1655, including the existence of a Pareto tradeoff between biomass synthesis and the capacity to generate maintenance energy. At a protein crowding of *f_MSA_* = 0.07, the predicted *μ_max_* of 0.95 h^-1^ correlated closely with experimental *μ_max_* of 0.97 ± 0.06 h^-1^ (red shaded area, Fig. 6C, supplementary data S7). Both strains had similar predicted protein crowding at their respective *μ_max_* (*f_MSA_* = 0.07) despite a 40% difference in maximum growth rate and substantial differences in SA:V ratios as a function of growth rate (Fig. 2B). Interestingly, strains MG1655 and NCM3722 are predicted to have a similar potential for *q*_ATP_ at their respective *μ_max_* (37 ± 5 and 37 ± 4 mmol ATP (g cdw h)^-1^) despite the faster growth rate of NCM3722 (Fig. 2A).

#### Testing prediction 2

Altering the applied NCM3722 SA:V ratio while holding *f_MSA_*= 0.07 constant caused the predicted *μ_max_* to deviate from the experimental value. The experimental SA:V was varied from +25% to −25% (Fig. 2B), and the *μ* vs. *q_ATP_* analysis was repeated (Fig. 6D, supplementary data S7). Additionally, the experimental MG1655 SA:V ratio was applied to the NCM3722 simulations (Fig 6D). The predicted *μ_max_* changed in a predictable manner; increasing SA:V predicted faster growth before the Pareto tradeoff occurred. Applying the MG1655 SA:V ratio to the NCM3722 fluxes predicted a *μ_max_* of 0.75 h^-1^, a substantial deviation from the experimental *μ_max_* of 0.97 h^-1^.

#### Testing prediction 3

*sMSAc-FBA* predictions of NCM3722 P/O numbers were quantitatively consistent with experimental numbers when experimental NCM3722 SA:V values were applied (Fig. 6E, supplementary data S7). Analogous to the MG1655 analysis (Fig. 5E), the predictions suggest the submaximal P/O numbers are actually optimal uses of the constrained *sMSA*. Predicted P/O numbers varied substantially when the applied SA:V was increased or decreased (Fig. 6E). Applying the experimental MG1655 SA:V ratios with experimental NCM3722 fluxes did not accurately predict strain phenotype (Fig. 6E, supplementary data S7). The predicted P/O number increased with decreasing growth rate, again suggesting *sMSA* is a considerable constraint only at high growth rates.

#### Testing prediction 4

The inflection point in the NCM3722 P/O number vs. *μ* relationship predicted the onset of overflow metabolism. The analysis predicted acetate overflow at *μ* = 0.68 h^-1^, which correlated reasonably well with the experimental onset at 0.75 ± 0.05 h^-1^. The P/O number vs. *μ* trends for seven different SA:V trends were considered (Fig. 6E, supplementary data S7). The trends demonstrate inflection points that were identified by taking the second derivative of the empirical polynomial fits. Altering the SA:V altered the predicted onset in a predictable manner. A summary of the predicted maximum growth rates and acetate overflow metabolism is presented in supplementary data S7.

#### Testing prediction 5

Experimental NCM3722 biomass yields on glucose were plotted as a function of growth rate (Fig. 6E, supplementary data S1, S7). Analogous to MG1655 data, the maximal experimental biomass yield corresponded closely with the inflection point for the P/O vs. *μ* relationship (0.66 vs. 0.68 h^-1^, respectively). Finally, it should be noted that the *sMSAc-FBA* predictions were remarkably accurate despite the majority of the NCM3722 data being measured using a recombinant strain with an altered membrane proteome.

#### Testing prediction 6

Basan *et al.* examined a recombinant *E. coli* NCM3722 strain which had a titratable SA:V ratio based on the overexpression of a ‘useless’, *lacZ* protein (70). While this was not the original intent of the study, the data was useful for testing the *sMSA* theory. High expression levels of *lacZ* caused the cell size to increase. The published cell dimensions were converted into SA:V ratios assuming a rod-shaped cell (supplementary data S2). The recombinant strain was grown in a fixed nutrient environment (MOPS + glucose) while the SA:V changes were modulated through titratable protein overexpression. *E. coli* cultures with larger total cell volumes, and therefore larger total surface areas, had slower growth rates due to smaller SA:V ratios (Fig. 6F). The mechanism of cell SA:V manipulation used in Basan *et al*. (70) differed from the mechanism used in Si *et al.* (2). However, the changes in growth rate with SA:V were consistent across studies and were consistent with the presented *sMSA* theory which proposes a positive correlation between membrane catalytic potential per g cdw (positive function of cellular SA:V and *f_MSA_*) and growth rate. The *sMSA* theory’s sensitivity to SA:V was also tested in prediction 2.

## DISCUSSION

The rules of bacterial cell design remain incomplete despite decades of research. For example, the design rules that link cell geometry and membrane protein capacity with central metabolism have not been defined previously on a quantitative basis for different strains, growing at different rates, with different SA:V ratios, and with different membrane crowding. Bacteria display exquisite control over cellular geometry and membrane properties which are proposed here to be biologically significant as these properties are major mediators of possible phenotypes (1, 3, 17, 19). *E. coli* cell length and diameter increase with growth rate, reducing the SA:V ratio (Fig. 2B, 2C) (2, 12). However, as shown here, the reduction in enzymatic potential from decreases in SA:V is offset by an increase in the capacity of the membrane to host proteins (Fig 3A, supplementary data S6). This property is hypothesized to be critical to competitive phenotypes; if membrane protein content were not essential to fitness, this protein hosting would likely remain independent of, or decrease with, increasing growth rate (19). This membrane property was integrated into a new, molecular level theory which combines metabolism, cell geometry, and membrane protein crowding. *sMSA* theory was used to predict maximum growth rates, acetate overflow metabolism, electron transport chain efficiency, and maintenance energy fluxes (6, 24, 46). The predictions were compared to experimental phenomics data, membrane proteomics data, and recombinant strains with titrated SA:V ratios (Fig. 5 and 6). In all cases, the theory was remarkably accurate without accounting for either cytosolic volume or cytosolic macromolecular crowding (6, 12, 24). The presented *sMSA* theory is flexible and can be readily implemented with numerous systems biology approaches including biochemical pathway analysis (*e.g.* elementary flux mode analysis) (71), resource balance analyses (72, 73), growth law development (37), and other FBA-based approaches (46). Our research efforts have started examining a number of these exciting avenues.

Much effort has documented the importance of volume-associated proteome investment and intracellular molecular crowding on phenotype (6, 12, 24). Volume-based processes are certainly central to phenotype (5, 6, 9), but we hypothesize that both surface area- and volume-associated processes are critical and concurrently influence phenotype (1). Surface area requirements for membrane associated enzymes do not correlate well with volume proxies like enzyme mass (Fig. 1C, supplementary data S1); membrane surface area requirements for enzymes are instead proposed to be distinct biophysical properties which impose unique constraints (17, 24, 46). Tight regulation of cellular geometry and therefore SA:V ratio would arguably be futile if surface area-associated substrate transport and volume-associated protein synthesis were not also tightly regulated. If a single geometric aspect were limiting, evolution could select cells with changes in either cell length or cell diameter to overcome the surface area or volume limitation, respectively. A potential strategy to overcome the limits of geometry is to evolve membranes with higher protein hosting capacities. Mitochondria are eukaryotic organelles specialized in ATP generation (12). While the geometries of mitochondria can be similar to bacteria, their membranes can host proteins at areal densities approximately two-fold higher than an *E. coli* membrane (24 ± 5.6 % vs. 40-50%) (4, 12, 74) (reviewed in supplementary data S3) (Fig. 7A). Mitochondria membranes have likely evolved the capacity to host high protein fractions while maintaining membrane integrity because they persist in the relatively constant environment of the buffered and relatively safe eukaryotic cytosol. The role of cell geometries in different bacterial morphologies, *e.g.* bacilli vs. cocci, or the membrane protein capacities of Gram-negative vs. Gram-positive membrane architectures are open questions that can be addressed using *sMSAc-FBA* (12, 75).

**Figure 7.**
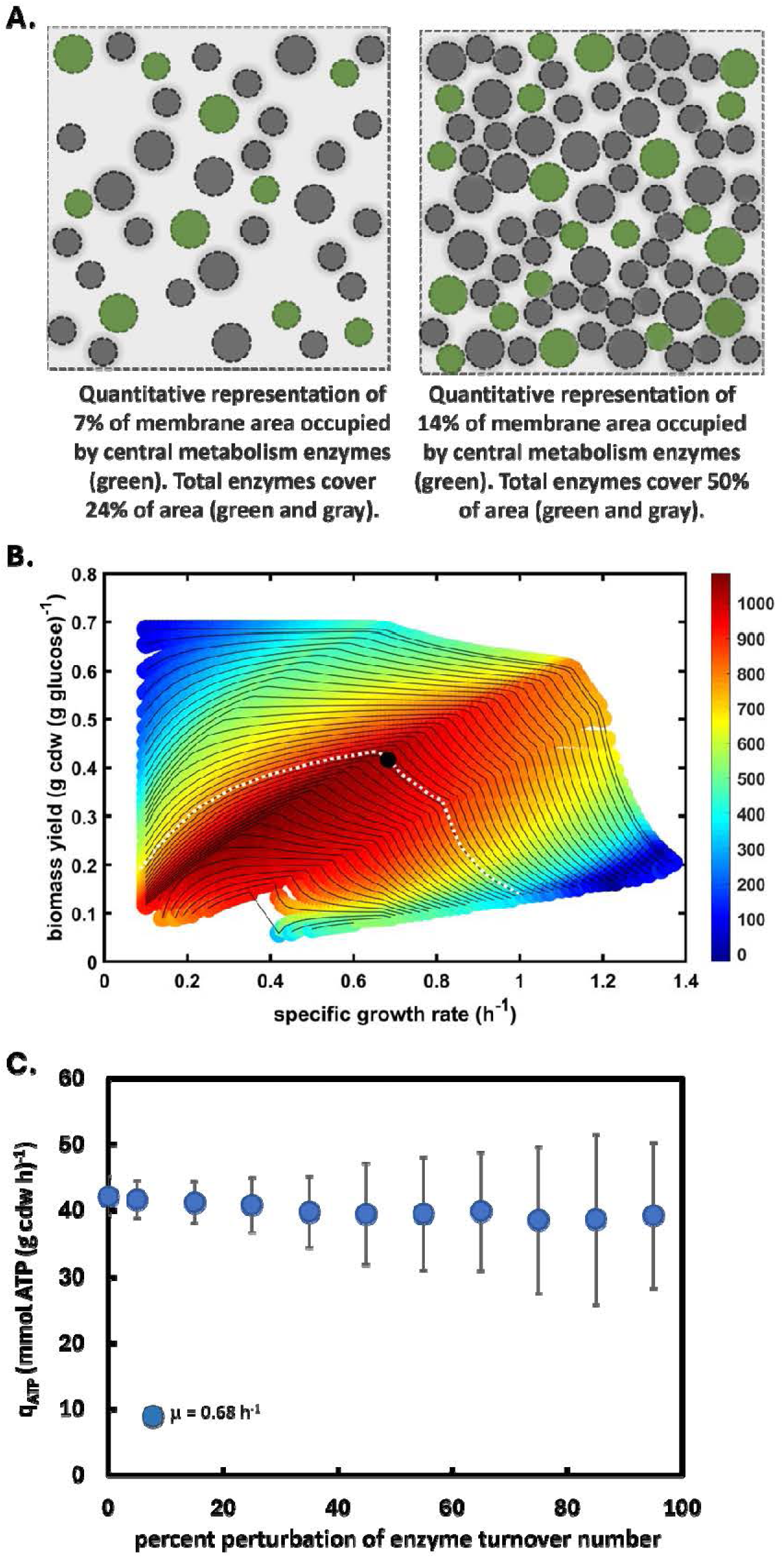
*sMSA* theory topics and implications. **A)** Quantitative representation of 24% and 50% of the membrane surface area being occupied by proteins. Areas are representative of bacteria *E. coli* and eukaryotic mitochondria inner membranes. **B)** Illustration of MG1655 experimental growth rate and biomass yield plotted in *sMSA* constrained phenotypic space. Dotted white line is a maintenance energy isocline. Third axis quantifies the density of ATP synthase complexes per membrane area. Experimental data suggests *E. coli* maximizes biomass yield for a specific maintenance energy requirement. **C)** Perturbation analysis of enzyme parameters and their effect on the potential for maintenance energy (*q_ATP_*) production. Experimental fluxes from batch growth of *E. coli* MG1655 on glucose salts media with *f*_MSA_ = 0.07. Enzyme parameters were independently perturbed within ±95% (abscissa) while experimental fluxes were simultaneously and independently perturbed within ± one standard deviation. Each point is the mean of 100 simulations, and error bars represent one standard deviation. *f_MSA_* = fraction of membrane surface area. *sMSA* = specific membrane surface area (nm^2^ gcdw^-1^).

*sMSAc-FBA* predictions assumed an idealized scenario including 1) all enzymes (except PtsG in MG1655 simulations) were saturated for substrate including ETC components and O_2_ cytochromes, 2) cells produced optimal levels of each enzyme with no overproduction, 3) all protein complexes were perfectly assembled with exact subunit ratios, 4) all enzymes were flawlessly embedded in the membrane, 5) no allosteric regulation of enzyme activity occurred based on membrane curvature (3, 76), and 6) the enzyme *k*_cat_ numbers increased two-fold for every 10°C increase in temperature (Q_10_ = 2) (31). The curated proteomics data provided a basis for estimating deviations from this idealized scenario.

Using a protein offset (Δ, Box 1) of ∼3% quantitatively aligns the *sMSAc-FBA* simulations with the experimental proteomics data without altering any presented interpretations or conclusions. The presented study implicitly included a perturbation analysis of Δ. For example, perturbing Δ for batch growth can be analyzed by examining results for *f*_MSA_ values larger or smaller than 0.07 (Fig. 5D, 6C).

Computational biology often assumes cells will adopt phenotypes that tend toward a maximum growth rate. This *in silico* growth rate is actually a relative maximum growth rate and not an absolute maximum growth rate. Fig. 7B plots the experimental MG1655 batch growth rate and biomass yield relative to the possible phenotypes based on the presented *sMSA* theory. The experimental data maps to a maximized biomass yield for a fixed maintenance energy demand, not a maximized growth rate. If the cell reduces maintenance energy demand, it could theoretically increase its growth rate while maintaining the same yield. For example, reducing the *q_ATP_* requirement by 8-10 mmol ATP (g cdw h)^-1^ could theoretically increase growth rate by 0.1 h^-1^ (shift experimental data point right +0.1 h^-1^ while keeping the same biomass yield). This proposed scenario, where a cell grows faster due to a decrease in the area associated with respiration activity (smaller *q_ATP_*) and increase in the area associated with substrate transporters, is consistent with some reported recombinant strain phenotypes (17, 77). Additionally, adaptive laboratory evolution can be used to create *E. coli* strains that grow faster than wild type cells, and some gene deletions can also increase growth rate (78, 79). Minimizing maintenance energy demand through reductions in futile cycling may contribute to the phenotypes. These strains increase growth rates, but likely at the expense of robustness. The presented work provides a quantitative framework to assess the changes.

Reported enzyme *k*_cat_ numbers vary substantially between published studies (supplementary data S1) (80). The presented study performed a literature survey of *k*_cat_ numbers and applied the same median *k*_cat_ values to all simulations for MG1655 and NCM3722 (supplementary data S3). The robustness of the presented results was analyzed by perturbing the *MRE*_i_ parameter which accounted for enzyme properties including *k*_cat_, degree of saturation, and surface area requirements (Box 1). The *MRE*_i_ parameter was randomly and independently perturbed within ±95% of the base value. Experimental fluxes were also perturbed randomly and independently within a standard deviation of the measured flux. Each scenario was analyzed with 100 simulations (Fig. 7C) (supplementary data S9). Capacity for *q_ATP_* was robust to perturbations; calculated *q*_ATP_ varied ≤ 10% for random perturbations of all *MRE*_i_ parameters up to ±95% of the base value (Fig. 7C) (supplementary data S9). *sMSA* theory is robust to the presented assumptions, likely based on the compensating nature of the redundant and parallel structures in metabolic networks (81). Given the robustness of the results, it is probable the parameters can be used in other animalcules with reasonable accuracy.

Additional robustness analyses examined perturbing one *MRE*_i_ parameter at a time to assess the effect on *q_ATP_*. These analyses considered five membrane-associated enzymes which accounted for >80% of the predicted surface area during batch growth (ATP synthase, glucose transporter (PtsG), ammonium transporter (AmtB), NADH dehydrogenase (Nuo), and cytochrome oxidase (Cyo); supplementary data S9). Each enzyme *MRE*_i_ was perturbed ±35% to increase or decrease the *sMSA* requirements, respectively. ATP synthase accounted for 45% of the surface area utilized by central metabolism enzymes and was most sensitive to *MRE*_i_ perturbations. Increasing the ATP synthase *MRE*_i_ parameter 35% while holding all other *MRE*_i_ parameters constant decreased the potential for *q_ATP_* fluxes by ∼26%, whereas decreasing the *MRE*_i_ parameter 35% increased the potential for *q_ATP_* fluxes by ∼8%. Perturbing the *MRE*_i_ parameter for the other four enzymes individually ±35% did not change the capacity for *q_ATP_* fluxes more than ±5% (supplementary data S9).

Biological optimality principles are of keen interest as they illuminate the (pseudo) rules of life and are readily implemented using systems biology methods. The presented study demonstrates that the commonly observed submaximal P/O numbers are likely not suboptimal strategies. Instead, they represent an optimal tradeoff between surface area requirements for substrate transporters and ETC components. Other commonly observed submaximal, yet likely optimized, phenotypes include overflow metabolisms occurring at both high (6, 82) and low growth rates (60), use of the Entner-Doudoroff (ED) glycolysis rather than Embden-Meyer-Parnas (EMP) glycolysis (81), and use of a partial citric acid cycle under nutrient limitation (83). Numerous *in silico* studies have replicated the overflow and ED vs. EMP glycolysis phenotypes using a variety of criteria (6, 14–16, 46, 81, 84). It is probable that many of these criteria are concurrently relevant to cell geometry (24, 46). All the criteria explicitly or implicitly define tradeoffs between resource sufficiency and resource scarcity and can be viewed through a lens of cellular economics. The relative value of a limiting resource is higher than for resources present in excess. Therefore, fitness is increased when use of the limiting resource is optimized (32, 81). The presented *sMSA* theoretical framework quantifies a metabolic resource defined by cell geometry and membrane protein crowding and elucidates cell design rules which govern selection of competitive phenotypes.

## Data Availability

All MATLAB code, models, and supplementary data can be found at github.com/rosspcarlson/carlson2026_sMSAcFBA. Additional *sMSAc-FBA* analyses, code, and reviewer comments from previous submissions can be found at (85).

## Author Contributions

R.P.C. conceived project, performed theory synthesis, developed code, analyzed data, wrote document, T.G., M.G.B., W.R.H., and R.M. analyzed data, developed interpretation, read and edited document, A.E.B. conceived project, developed code, analyzed data, read and edited document.

## ACKNOWLEDGEMENTS

R.P.C. would like to acknowledge Kristopher Hunt, Thomas Cronimus, Adrienne and Arnold for insightful discussions and feedback. R.P.C. would like to acknowledge Terrance Hwa for sustained encouragement.

R.P.C. was supported in part by NIH award U01EB019416 and AFOSR award FA9550-23-1-0589. A.E.B. was supported in part by USDA National Institute of Food and Agriculture grant no. 2021-67013-34537 and NSF grant no. DEB 2238670.

## Summary of Supplementary Materials

Files available at: github.com/rosspcarlson/carlson2026_sMSAcFBA

**Supplementary Data S1:** Literature survey of extracellular flux data for E. coli K-12 strains MG1655 and NCM3722 including consensus data sets. Comparison of enzyme volume and membrane surface area requirements for 22 central metabolism enzymes.

**Supplementary Data S2:** Literature values for cell geometry as a function of growth rate for *E. coli* K-12 strains MG1655 and NCM3722. MreB and FtsZ titration data for MG1655. Cell volume data for titration of cell ‘inflating’.

**Supplementary Data S3:** Literature review of enzyme and biophysical parameters used in study.

**Supplementary Data S4:** *in silico,* stoichiometric model and associated data files used in the presented study to represent *E. coli* strains MG1655 and NCM3722.

**Supplementary Data S5:** Summary of MATLAB code used for each figure in study and Github information.

**Supplementary Data S6:** *E. coli* K**-**12 membrane proteomics data from three independent studies collated by Belliveau et al. 2021.

**Supplementary Data S7:** Summary of simulation output for *E. coli* K**-**12 strains MG1655 and NCM3722 constrained by consensus fluxes, SA:V, *f_MSA_*, and when appropriate PtsG saturation.

**Supplementary Data S8:** Summary of experimental and in silico P/O number calculations.

**Supplementary Data S9:** Analysis of maintenance energy fluxes as a function of growth rate, membrane surface area, protein hosting capacity, and P/O number. Perturbation analysis of enzyme *MRE* parameters.

---

**Box 1.**
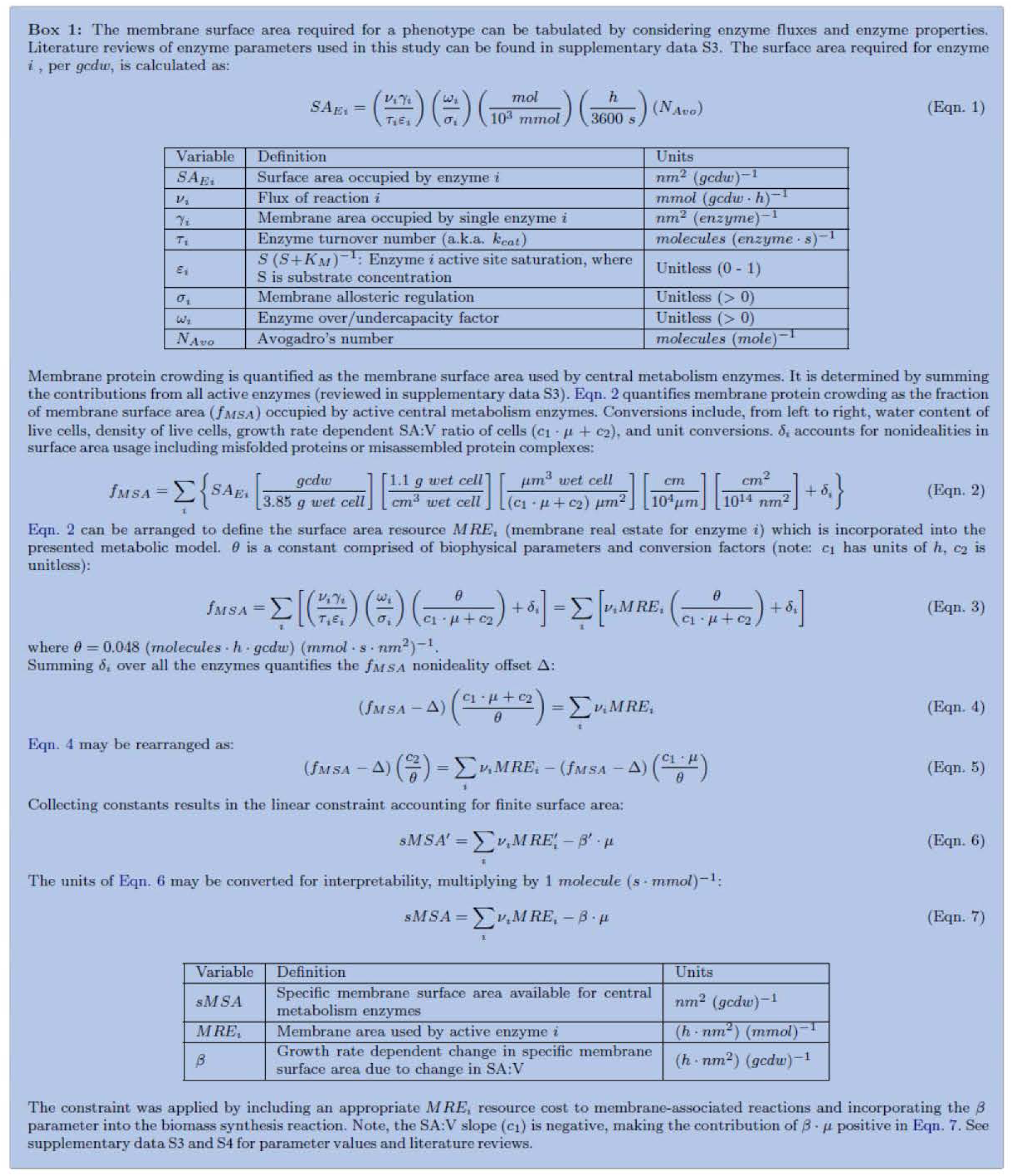

XXXXXXXXXXXXXXXXXXXXXXXXXXXXXXXXXXXXXXXXXXXXXXXXXXXXX XXXXXXXXXXXX

**Reviews and rebuttals from three submissions spanning March 2024 to November 2025**

R.P.C. would like to acknowledge and thank the reviewers for their investment of time.

**PLOS Computational Biology submitted September 5, 2025**

**reviews received November 10, 2025**

** On Behalf Of PLOS Computational Biology**

**Sent: Monday, November 10, 2025 10:44 AM To: Carlson, Ross**

**Subject: PLOS Computational Biology Decision: PCOMPBIOL-D-25-01810 - [EMID:e73d4019720e28fd]**

**\*\*External Sender\*\***

CC: “Tomáš Gedeon”, “Mauricio Garcia-Benitez”, “William R. Harcombe”, “Radhakrishnan Mahadevan”, “Ashley E. Beck”

**PCOMPBIOL-D-25-01810**

Cell Geometry and Membrane Protein Crowding Constrain Growth Rate, Overflow Metabolism, Respiration, and Maintenance Energy.

PLOS Computational Biology Dear Dr. Carlson,

Thank you for submitting your manuscript to PLOS Computational Biology. As with all papers, your manuscript was reviewed by members of the editorial board. Based on our assessment, we have decided that the work does not meet our criteria for publication and will therefore be rejected. If external reviews were secured, reviewers’ comments will be included at the bottom of this email.

We are sorry that we cannot be more positive on this occasion. We very much appreciate your wish to present your work in one of PLOS’s Open Access publications. Thank you for your support, and we hope that you will consider PLOS Computational Biology for other submissions in the future.

Yours sincerely,

Stefan Klumpp Academic Editor

PLOS Computational Biology

Marc Birtwistle Section Editor

PLOS Computational Biology

**Additional Editor Comments (if provided):**

As you can see the reviewers are divided in their assessment of your manuscript. One reviewer states that your paper is excellent, but still raises substantial points, while the other is much less enthusiastic and lists many experimental results inconsistent with your model. Some of the critical issues also appear rather similar to issues raised in the previous reports that were included with the submission.

[Note: HTML markup is below. Please do not edit.]

**Reviewers’ Comments (if peer reviewed):**

Reviewer’s Responses to Questions

**Comments to the Authors:**

**Please note here if the review is uploaded as an attachment.**

**REVIEWER 1 COMMENTS**

“Reviewer #1: Carlson et al. propose a metabolic theory to predict growth rate and overflow metabolism. They argue that membrane protein crowding limits growth rate and determines the onset of overflow metabolism in bacteria.

Evidence they provide includes the different growth rates, surface to volume ratios and rates of overflow metabolism of MG1655 and NCM3722 strains.

However, very similar models and mechanisms, sometimes referred to as the known as the membrane real estate hypothesis, have been proposed many times before (e.g. PMID: 28755958). The novelty of this work in the context of literature is modest.”

RESPONSE: Thank you for reviewing the manuscript, we appreciate your investment of time. Alternative viewpoints are always appreciated even if we disagree. Alternate viewpoints highlight a mutual need to keep an open mind.

Similar models have not been proposed ‘many times’; there are two publications. Our work is the third. The introduction of the document explicitly covers this critique. This is the third study and is by far the most comprehensive, quantitative, and predictive (please see Reviewer 2’s comments acknowledging this contribution). Coauthor Mahadevan proposed the first theory in 2011 (10.1038/msb.2011.34). The study was predictive but qualitative due to a lack of quantitative data in 2011. The second study (mentioned by Reviewer 1) was from 2017 and is quantitative but not predictive on a systems level, as it is comprised of equations in an Excel spreadsheet. The current study is predictive, quantitative, based on molecular level details, explicitly accounts for growth rate dependent changes in SA:V ratios, explicitly accounts for growth rate dependent membrane protein crowding, and compares two distinct *E. coli* strains with different phenotypes. It integrates data from more than a dozen physiological studies and three proteomics studies and proposes new design rules for bacterial cells. The current study quantitatively predicts maximum observed growth rate, respiration efficiency, and overflow metabolism onset from the intersection of membrane constraints and metabolism. This is all stated with appropriate citations in the introduction and represents a substantial advance in theory and practice.

“Moreover, the hypothesis and model predictions are in direct contradiction with various experimental findings. Therefore, I cannot recommend publication of this manuscript.”

RESPONSE: We find this statement to be inaccurate. The presented theory and the Reviewer’s points are not in direct contradiction. In fact, the Reviewer’s phenotypic examples are consistent with the presented theory. We have integrated the ‘supposedly contradictory’ experimental findings into the manuscript as additional evidence supporting our claims (examples described below).

Here are some experimental observations by the Hwa lab and others that seem to directly contradict the proposed model.

1. “Overexpression of useless bulk protein results in very large cells that grow very slowly. Metabolic flux requirements per biomass linearly decrease with growth rate. Cells have ample membrane surface for respiration, yet continue to ferment (PMID: 26632588, PMID: 26519362).”

RESPONSE: Thank you for highlighting this study. These experimental findings are consistent with the presented theory, and we have integrated the data into the manuscript as additional supporting evidence (tested prediction 6, page 10). Geometrical scaling dictates that cell volume increases faster than surface area (R^3 vs. R^2); therefore, large cells have smaller SA:V ratios relative to small cells. Consequently, large cells are often limited for surface area to support the substantial volume-based metabolic processes like the synthesis of unused proteins. Imbalances between surface area and volume can result in phenotypes like acetate overflow. Figure 5F presents experimental data for *E. coli* MG1655 showing a negative relationship between cell size and growth rate and also shows a positive relationship between SA:V and growth rate. Additionally, we have added experimental data from the Hwa group which shows the same SA:V vs. growth rate relationship with an *E. coli* NCM3722 derivative (Figure 6F).

Finally, Figure 6D quantifies the effect of NCM3722 SA:V ratios as applied by the presented theory. The experimental scenario raised by Reviewer 1 is consistent with and supportive of our proposed theory.

**Figure 5F.**
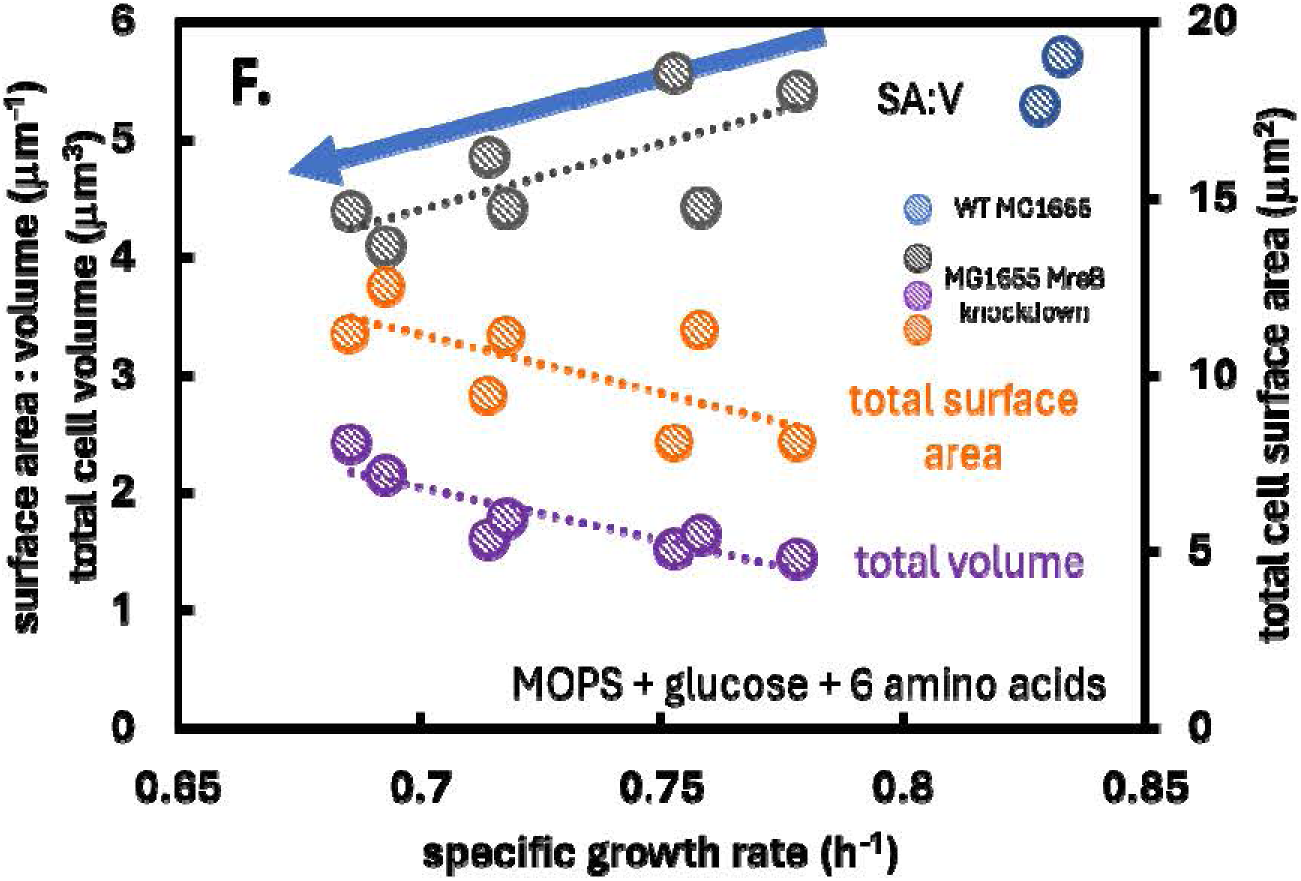
Experimental *E. coli* MG1655 data measuring the effect of titrated surface area to volume ratios (SA:V) and cell size (volume) on growth rate. Original data from 10.1016/j.cub.2017.03.022. Full caption can be found in manuscript.

**Figure 6F:**
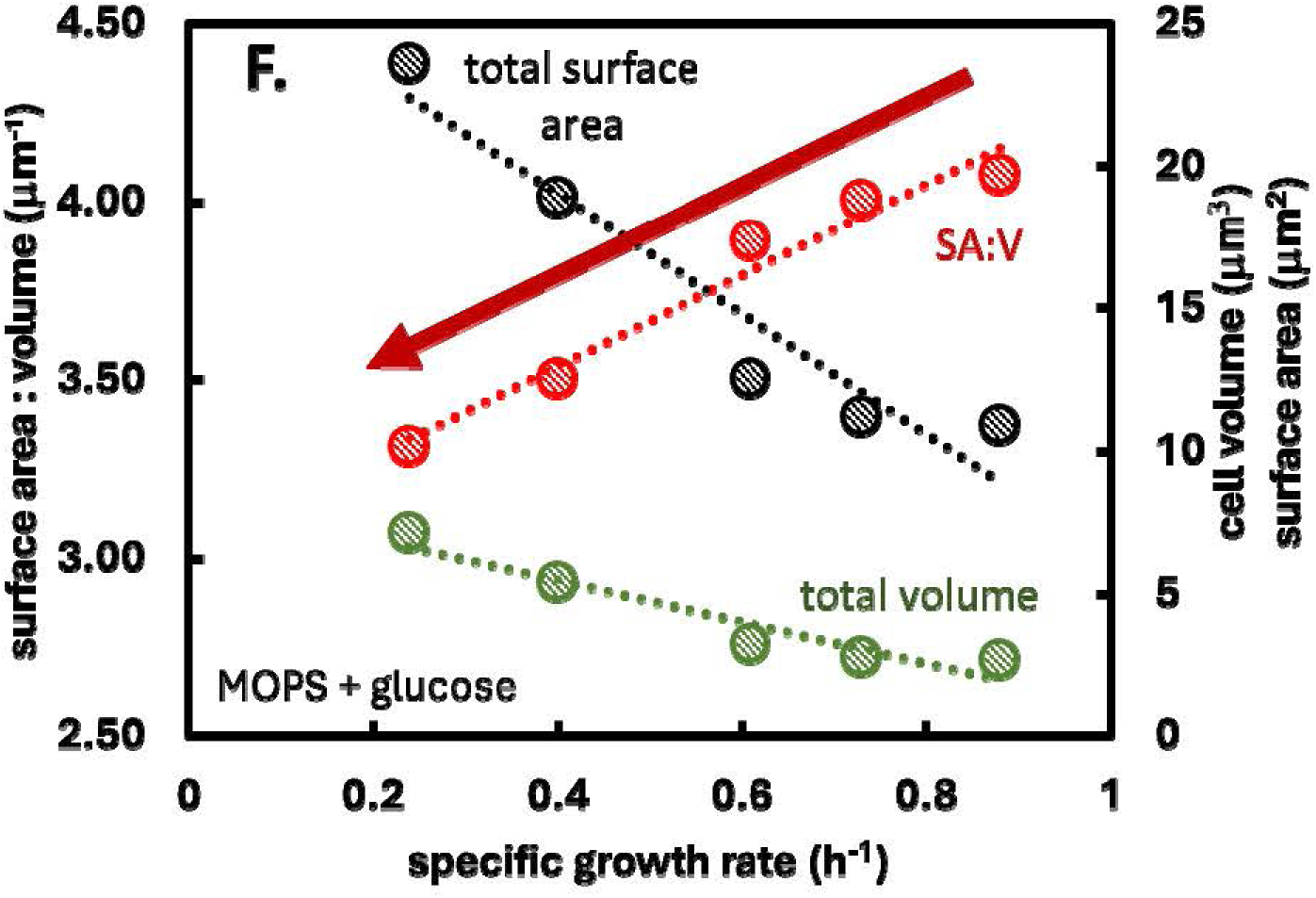
SA:V data plotted vs specific growth rate for strain NQ1389. The strain modulates SA:V ratios based on overexpression of a ‘useless’ protein. Data from Basan et al. 10.15252/msb.20156178. Full caption can be found in manuscript.

**Figure 6D.**
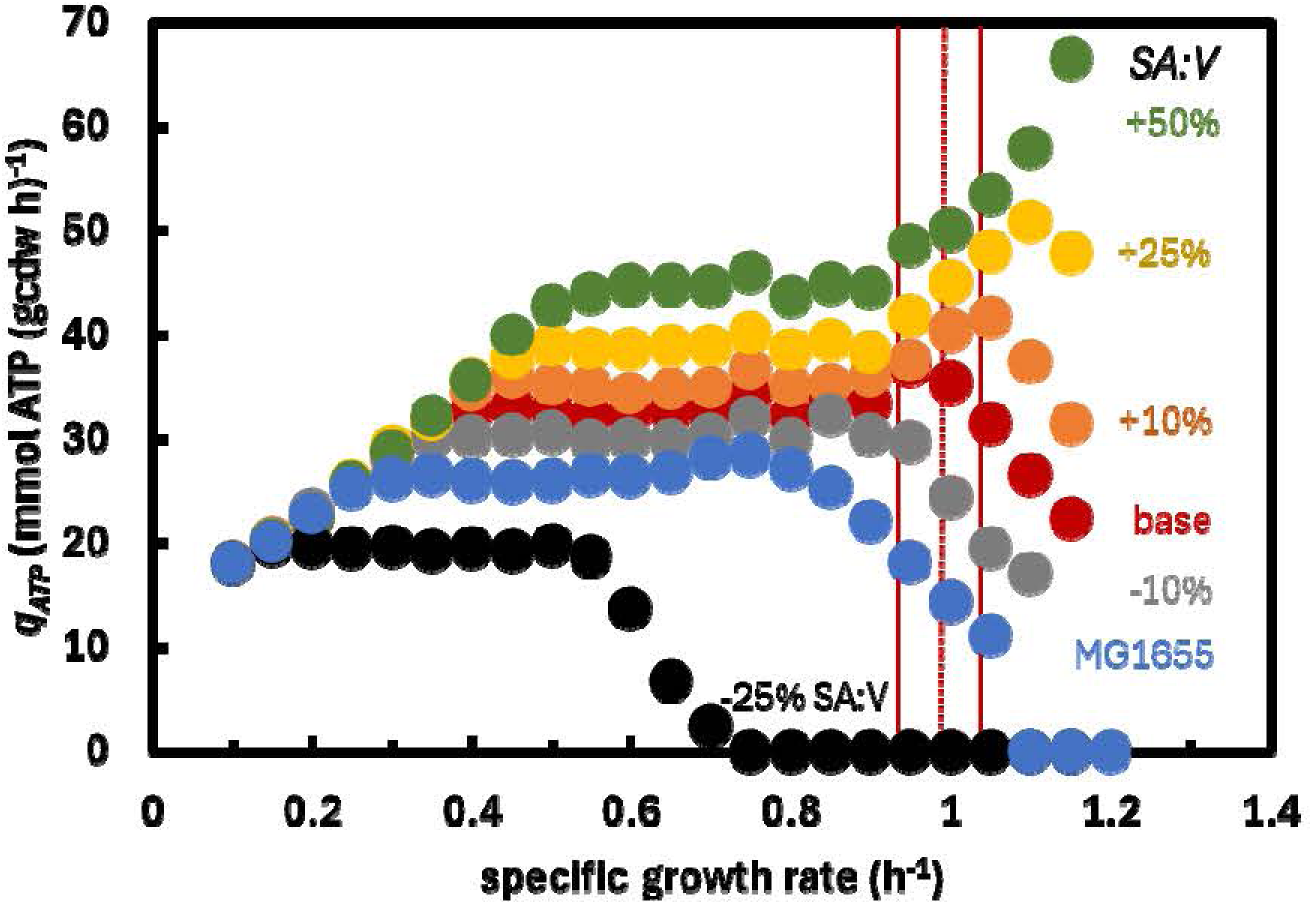
*E. coli* NCM3722 simulation data quantifying the effect of surface area to volume (SA:V) on the potential of experimentally measured fluxes to produce ATP. See text for details. Full caption can be found in manuscript.

2. “Knocking down respiration results in faster growth rates and higher overflow metabolism in many different conditions (PMID: 26519362). The opposite would be expected from the model proposed by Carlson et al. It should not be possible to speed up growth rate by forcing cells to ferment by reducing respiration.”

RESPONSE: The reviewer’s critiques are not consistent with the presented theory nor the presented data. The scenario proposed by Reviewer 1 is in fact plotted in Figure 4 (and the data is in supplementary material S6) and is therefore predicted by the theory. Please direct your attention to Figure 4A, a growth rate of 0.1/h and a yield of 0.35 g/g. At this growth rate, the cell has sufficient surface area, low glucose fluxes, high efficiency respiration, and presents no acetate overflow. Holding the yield constant, move to the right along the dotted red line: this quantifies the effects of increasing growth rate. As the growth rate increases, the glucose flux increases, and at a growth rate of ∼0.5/h acetate overflow initiates due to scarcity of surface area per volume. Continue moving right holding the yield constant, and glucose flux increases, overflow increases, and respiration decreases due to limited surface area per volume. This is exactly the scenario described by Reviewer 1. Reducing respiration-associated proteins frees up valuable membrane surface area for substrate transporters which could, under appropriate conditions, increase growth rate. Reducing respiration fluxes would inherently increase overflow fluxes to balance metabolic electrons. An example of this scenario related to gene deletions and adaptive laboratory evolution was added to the discussion, with appropriate citations, after the Reviewer raised the same comment during the last submission.

**Figure 4A.**
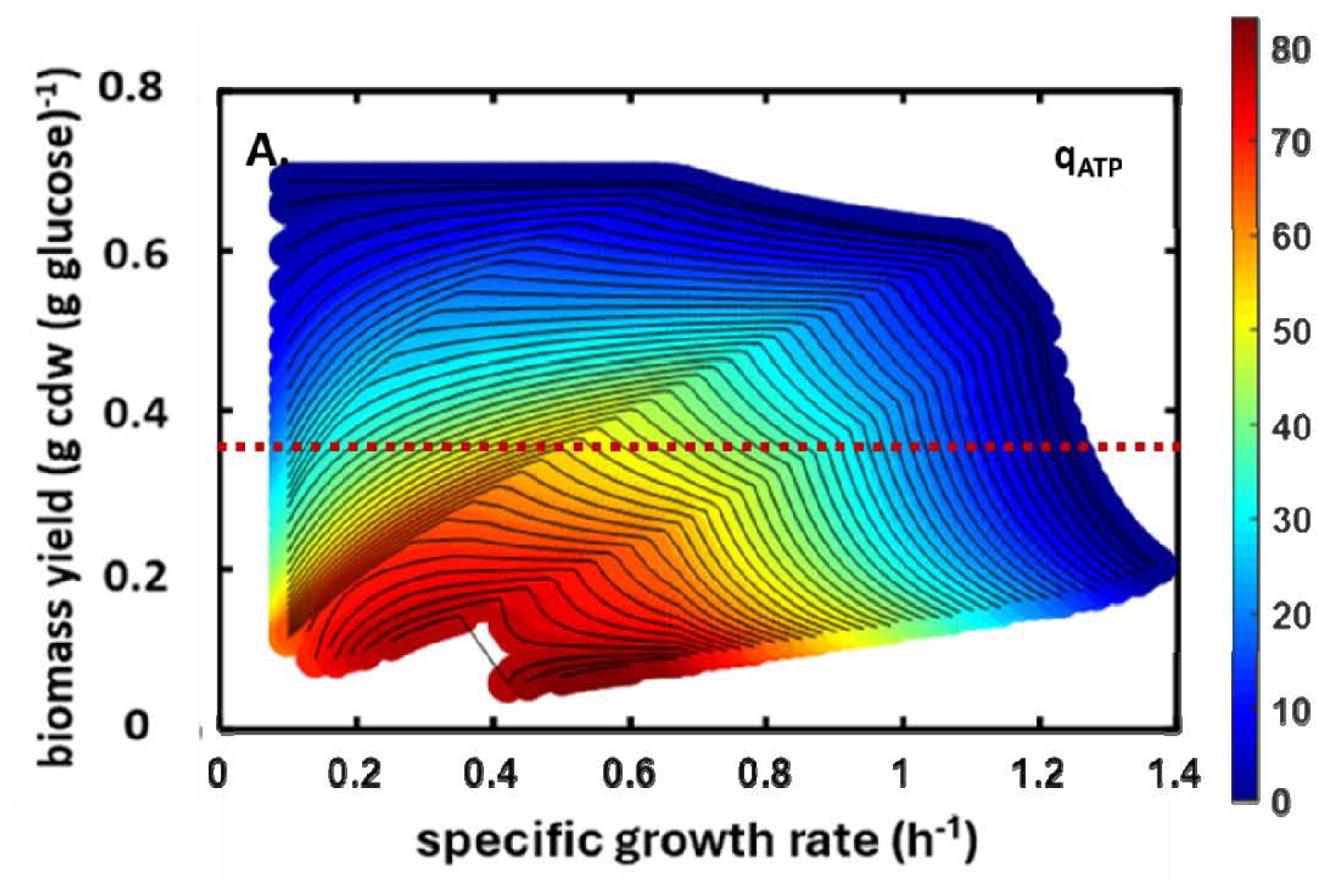
Simulation data for *E. coli* MG1655 demonstrating the scenario proposed by Reviewer 1. Simulation data can be found in Supplementary Data file S6. Full caption can be found in manuscript.

3. “Growth rates between different substrates can be swapped by exchanging promoters of uptake systems (PMID: 37662352). Cell size and surface to volume ratio appears to follow metabolism rather than the other way around. Growth rates do not appear optimal inn terms of growth rate maximization.”

RESPONSE: Our document does not claim cells attain their absolute maximum possible growth rate; in fact, we argue that cells do not maximize their absolute growth rate but instead attain a relative maximum growth rate and that cells can be evolved for higher growth rates. The text states “This computational growth rate is, in actuality, a relative maximum growth rate and not an absolute maximum growth rate. Fig. 7B plots the experimental MG1655 batch growth rate relative to the possible growth rates based on the presented *sMSA* theory. The experimental data maps to the maximization of biomass yield for a fixed maintenance energy demand not the maximization of absolute growth rate. If the cell reduces maintenance energy demand, it could theoretically increase its observed growth rate while maintaining the same yield.” Additionally, please see Figure 4A and associated text. Our document claims that cell growth rate increases until the finite membrane surface area results in a decrease in the capacity to generate maintenance energy (please see Figures 5C and D). This defines a Pareto tradeoff that predicts the observed maximum growth rate (observed maximum growth rate < absolute theoretical maximum growth rate). This is a key hypothesis developed using *E. coli* MG1655 data and tested using *E. coli* strain NCM3722 data (please see Figure 6C and associated text). Both MG1655 and NCM3722 utilize the available SA:V and protein crowding in the same manner enabling the prediction and explanation of phenotype from a calibration data set.

We agree with the reviewer’s postulation regarding cell dimensions and growth; the text also agrees with the Reviewer’s statement. Cell size and SA:V ratio are a function of growth rate which is a result of nutrient environment and Monod like growth laws. This is shown experimentally in Figure 2B. The document has a new sentence and three new references emphasizing this view on p. 3. This relationship is expected to hold regardless of swapped substrate promoters.

However, it is also possible to influence phenotype by modulating cell geometry with recombinant tools. This is shown in Figures 5F and 6F, where modulating the cellular SA:V ratio in a fixed nutrient environment alters the growth rate in a predictable manner.

**Figure 5C and 5D.**
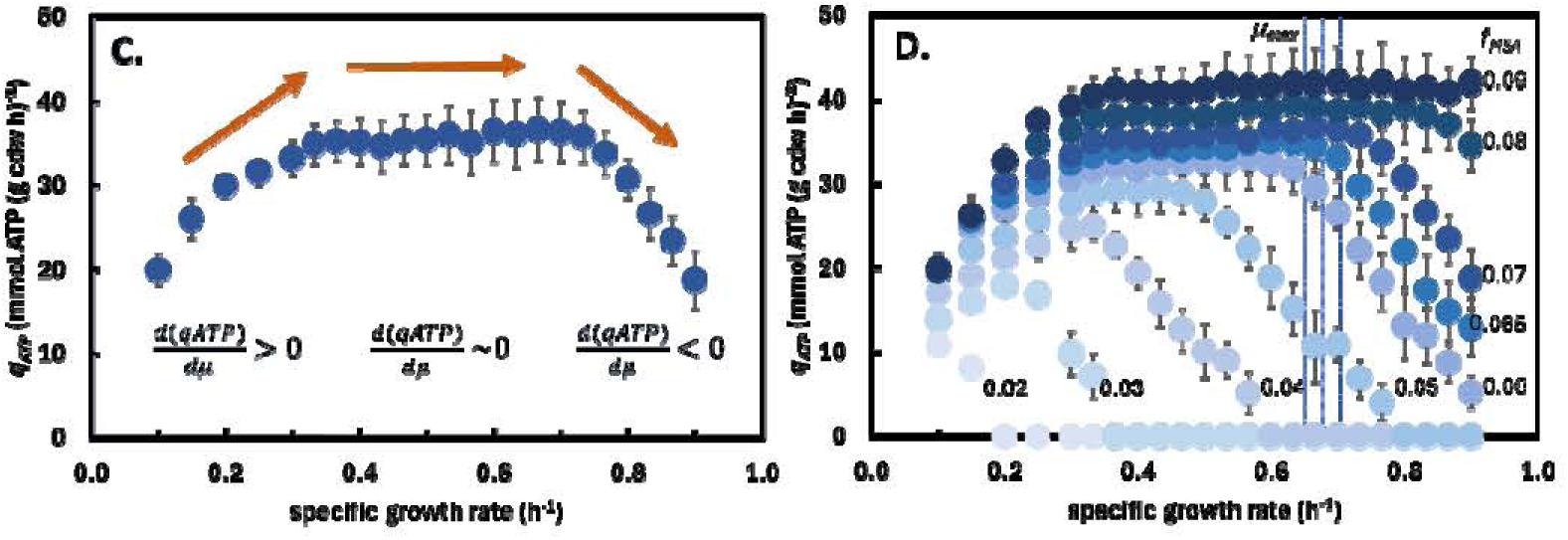
Pareto Front between growth rate and maintenance energy fluxes for strain *E. coli* MG1655. Simulations suggested the absolute maximum growth rate is 1.4/h while the experimentally observed maximum growth rate is ∼0.7/h consistent with our theory. Full caption can be found in manuscript.

**Figure 6C.**
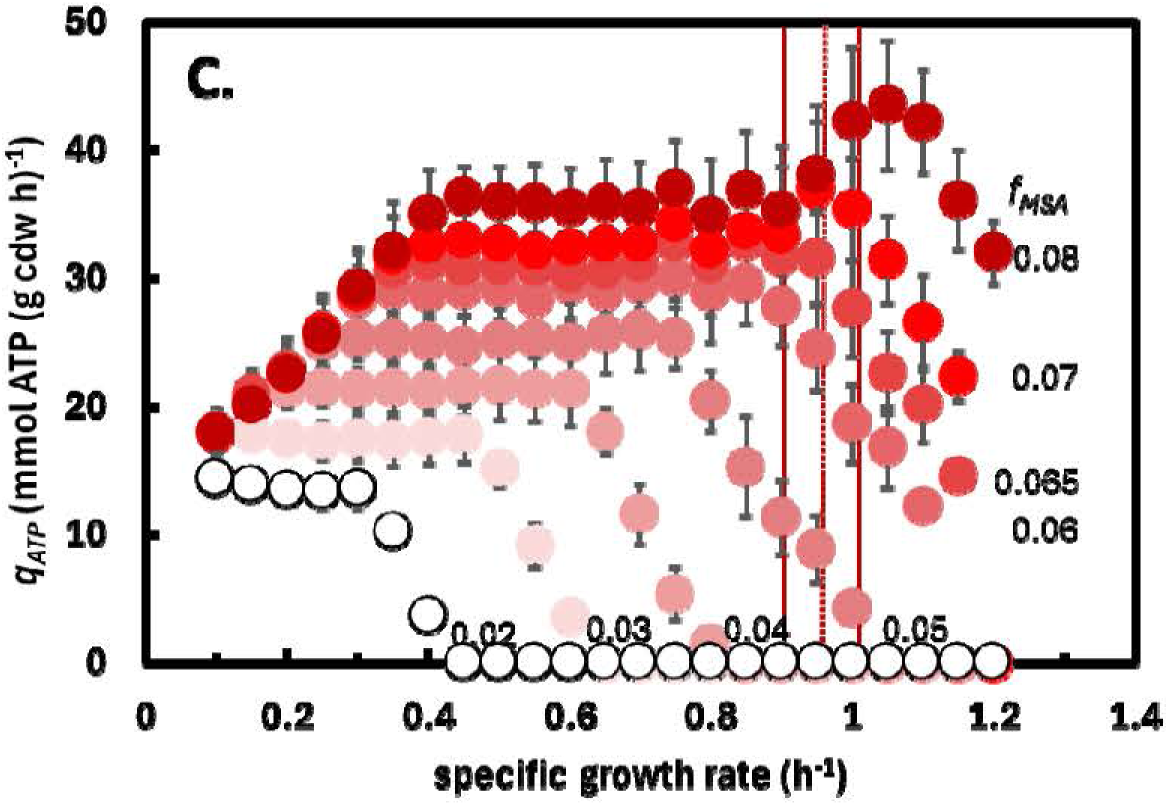
Pareto front between growth rate and maintenance energy fluxes for *E. coli* strain NCM3722. The simulations suggest the absolute maximum growth rate is >1.4/h while the experimentally observed maximum growth rate is ∼1/h consistent with our theory. Full caption can be found in manuscript.

**Figure 2B.**
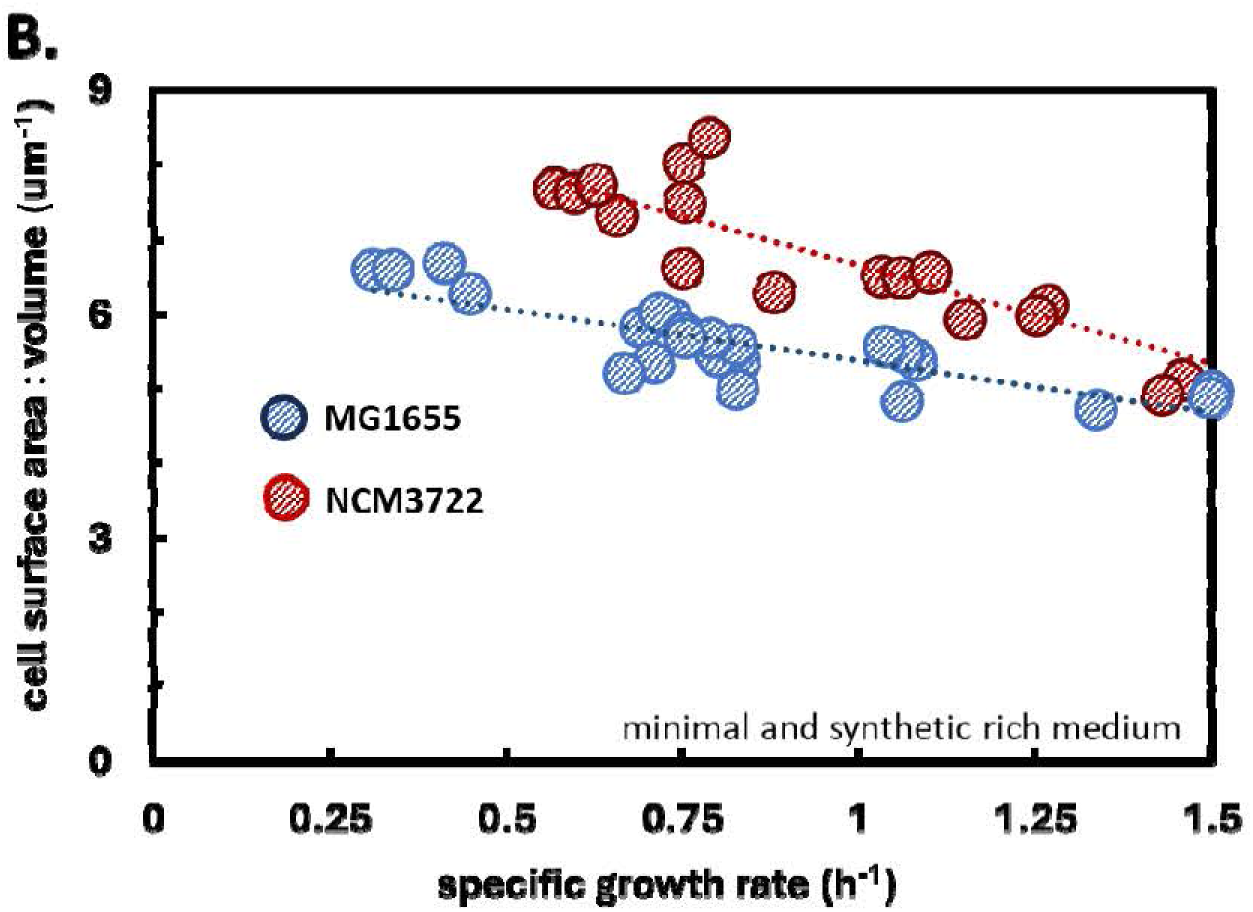
Experimental data showing relationship between *E. coli* cell surface area to volume and growth rate. Original data from 10.1016/j.cub.2017.03.022. Note the MG1655 data is for wild type strains acclimating to different nutrient environments by changing growth rate and cell dimensions. Full caption can be found in manuscript.

**Figure 5F.**
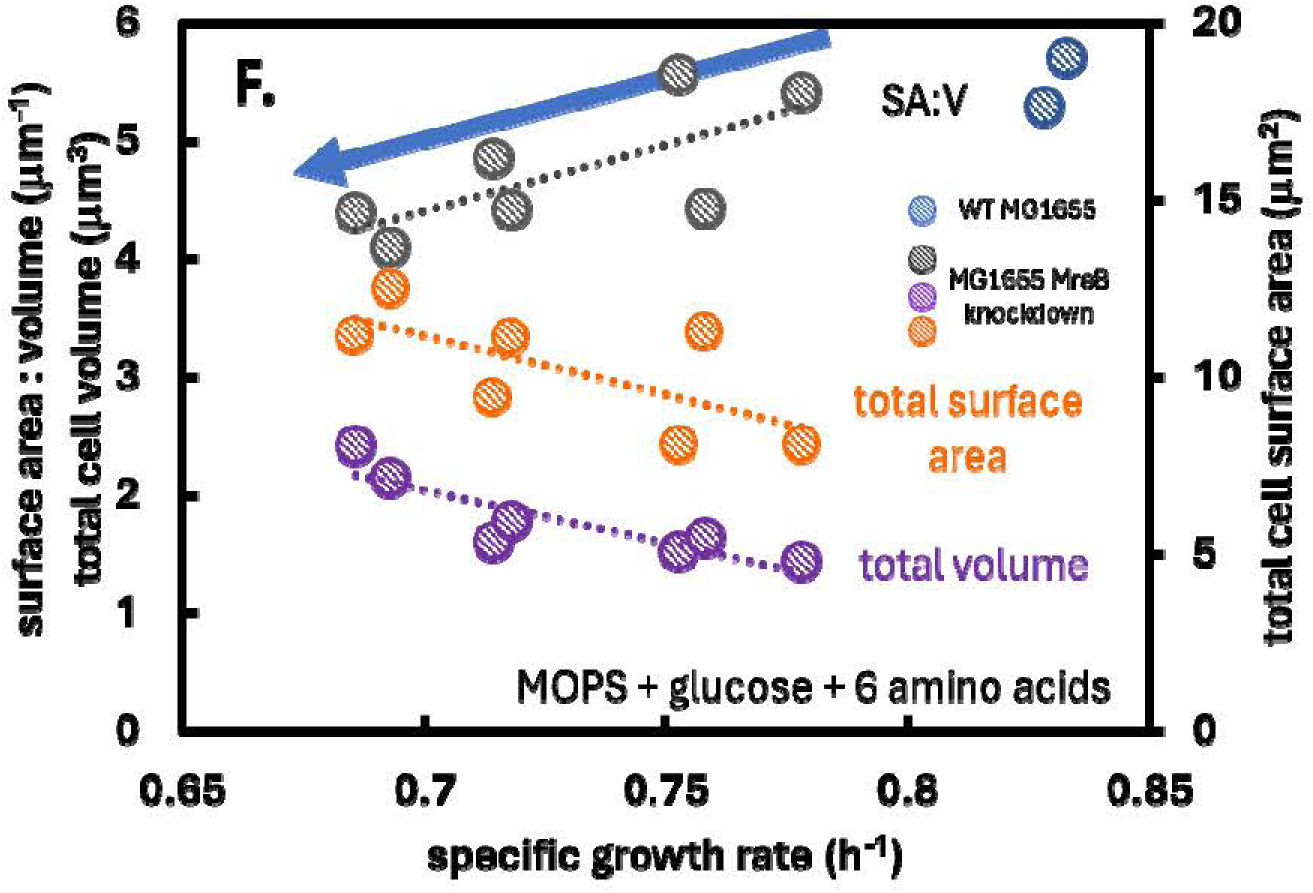
Experimental *E. coli* MG1655 data measuring the effect of titrated surface area to volume ratios (SA:V) and cell size (volume) on growth rate. Original data from 10.1016/j.cub.2017.03.022. Note, the SA:V ratios are modulated by changing the transcriptome while the nutrient environment is not changed. Full caption can be found in manuscript.

**Figure 6F:**
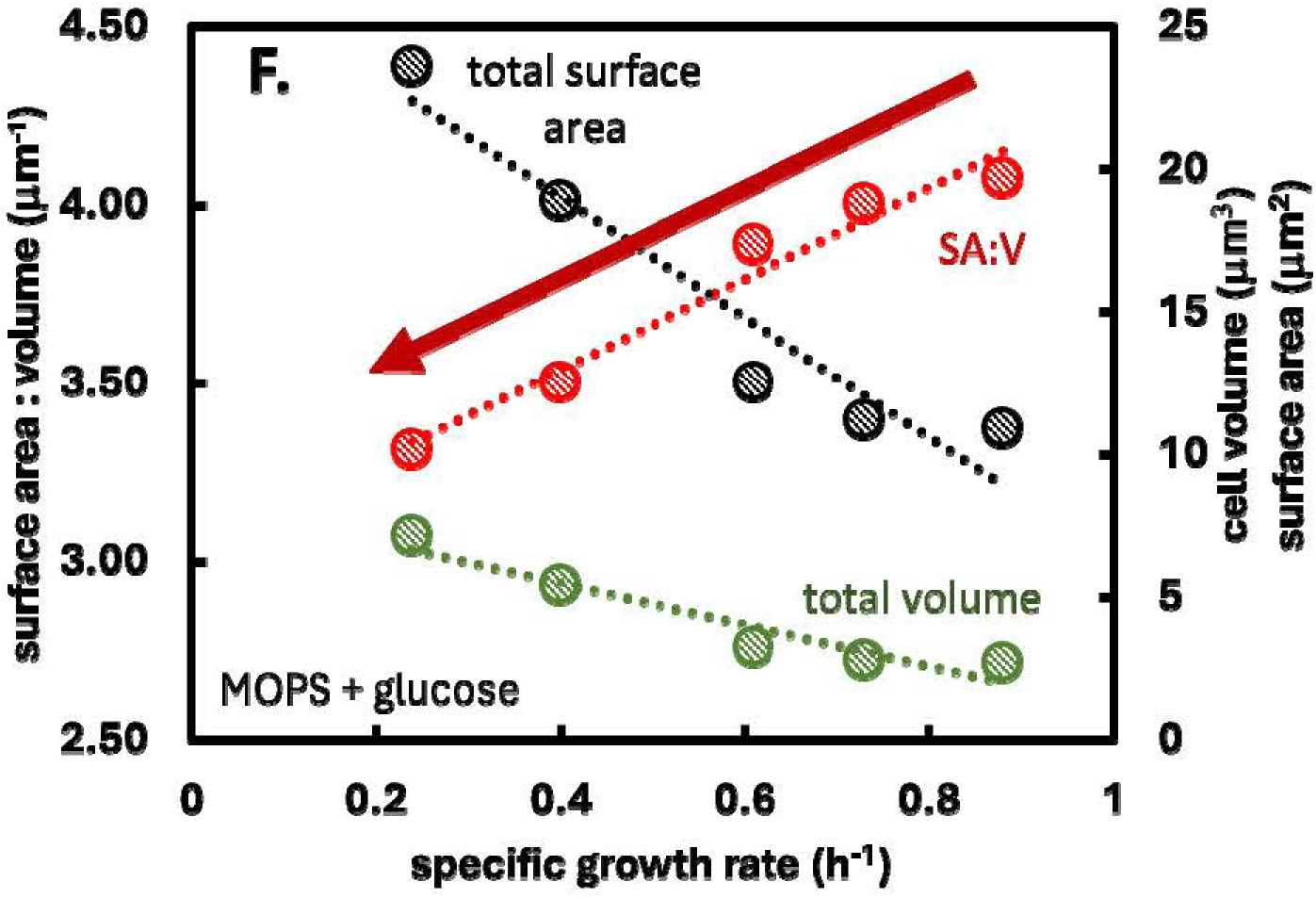
SA:V data plotted vs specific growth rate for strain NQ1389. The strain modulates SA:V ratios based on overexpression of a ‘useless’ protein. Data from Basan et al. 10.15252/msb.20156178. Full caption can be found in manuscript.

4. “Surface to volume ratio of bacteria can be changed by genetic titration of ftsZ. However, this has little to no effect of growth rates (PMID: 31104932). From the model proposed by Carlson et al., longer cells should suffer reduced growth defects, whereas shorter cells should have more capacity for respiration, enabling faster growth.”

RESPONSE: Thank you for the comment. Two separate studies titrated *E. coli* SA:V ratios and measured culture growth rates; the results are in accordance with sMSA theory (Figure 5F and Supplementary data S2). The Jun group at UCSD has published an exceptional dataset (10.1016/j.cub.2017.03.022). They built *E. coli* MG1655 strains with titratable FtsZ or MreB. Both proteins modulate cell dimensions and therefore SA:V. As stated in our document, titration of FtsZ did not result in substantial changes in cell SA:V ratios nor did it substantially change the strain growth rate. However, the MreB system achieved substantial changes in cell SA:V ratios. As the SA:V ratio decreased, the strain grew slower consistent with the presented theory. Additionally, the Hwa group engineered a strain to overproduce a useless protein to the extent that cell size changed. The larger cells had smaller SA:V ratios and grew slower consistent with the presented theory. Our revised document has expanded the number of predictions developed on *E. coli* MG1655 and tested on *E. coli* NCM3722 from 5 to 6 to include this example (page 10).

**Figure 5F.**
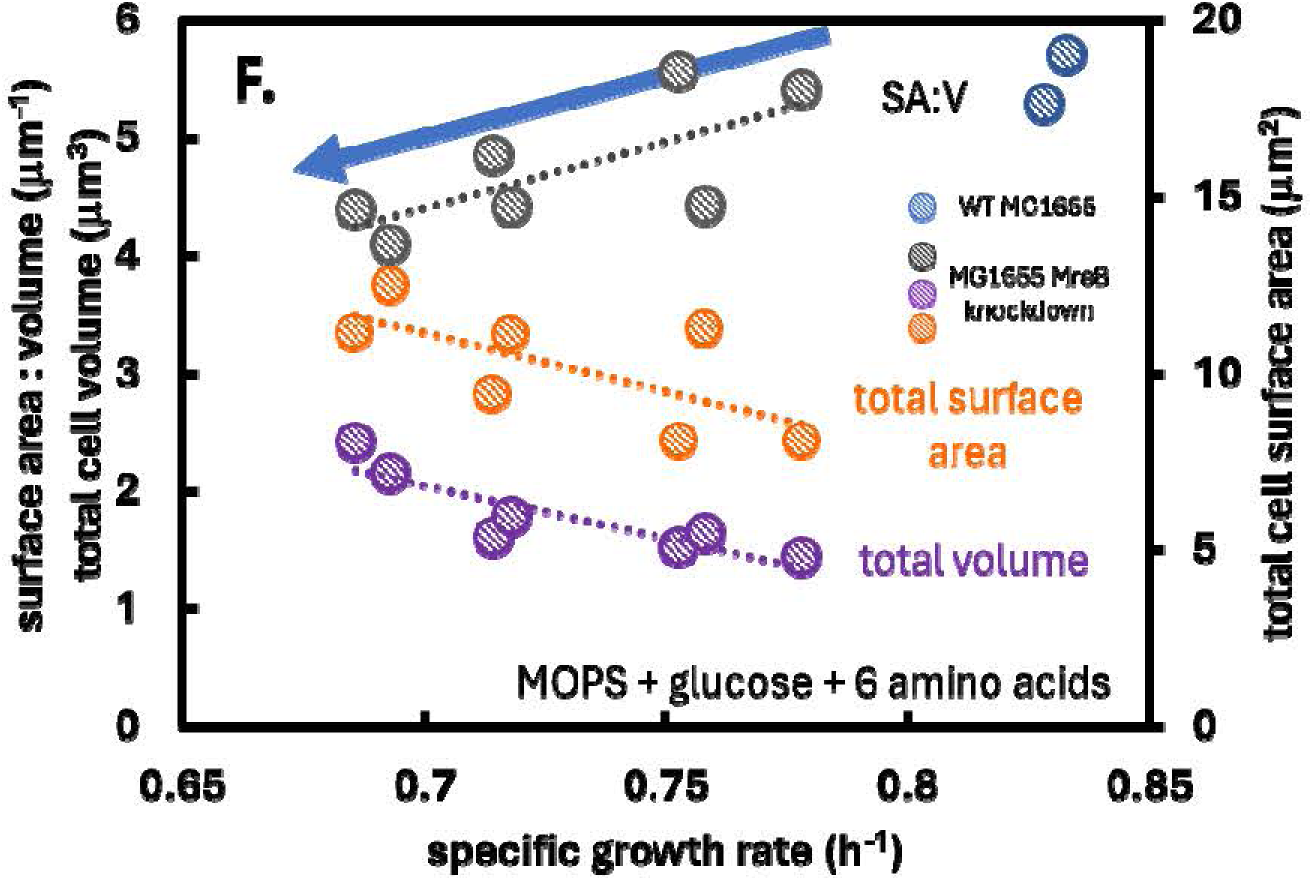
Experimental *E. coli* MG1655 data measuring the effect of titrated surface area to volume ratios (SA:V) and cell size (total volume) on growth rate. Original data from 10.1016/j.cub.2017.03.022. Full caption can be found in manuscript.

**Figure 6F:**
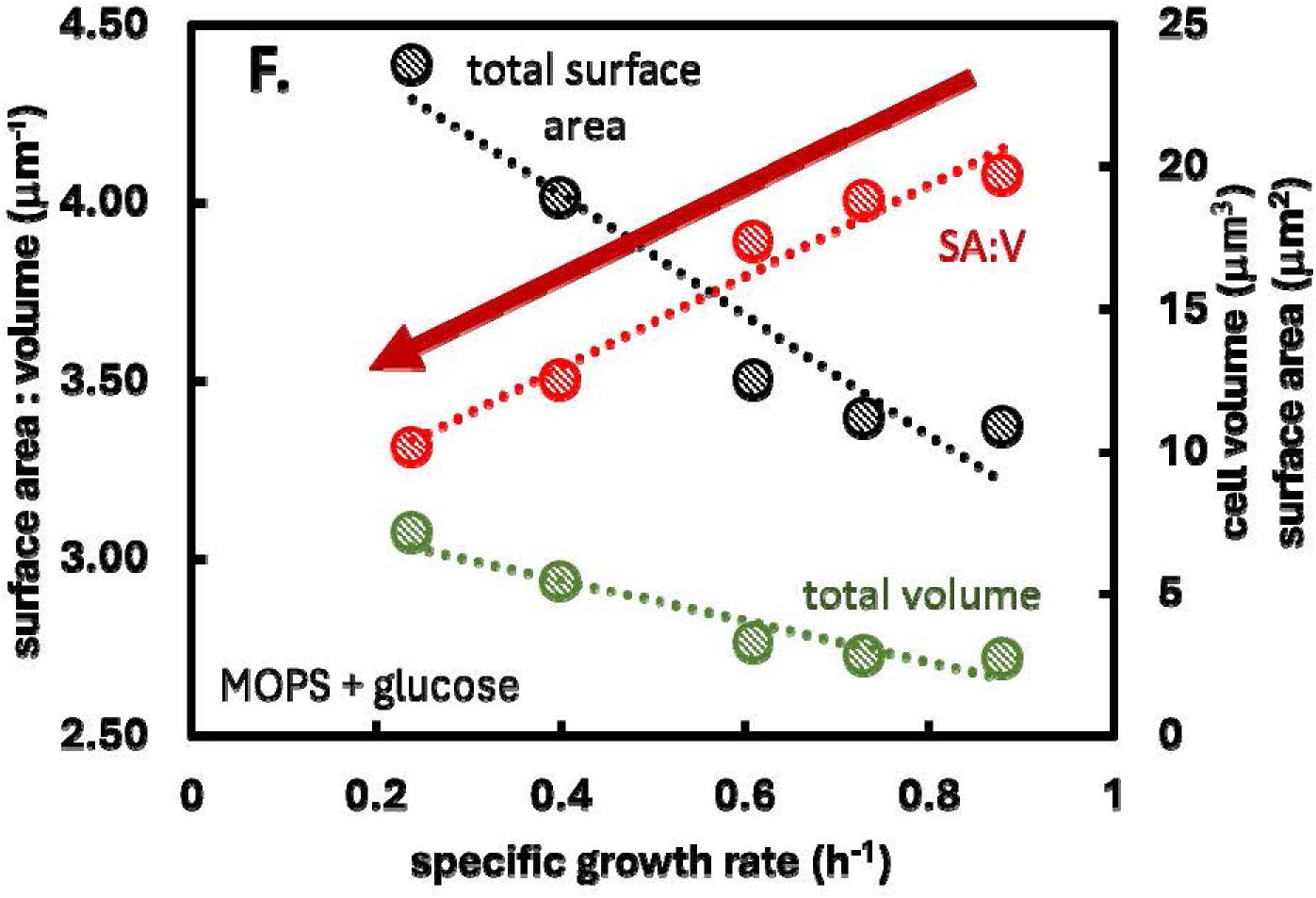
SA:V data plotted vs specific growth rate for *E. coli* strain NQ1389. The strain modulates SA:V ratios based on overexpression of a ‘useless’ protein. Data from Basan et al. 10.15252/msb.20156178. Full caption can be found in manuscript.

**REVIEWER 2**

Reviewer #2: In their manuscript, the authors present a theoretical framework that links cell geometry and membrane protein crowding with key phenotypic properties of microbial metabolism, such as growth rate, respiration efficiency and overflow metabolism. They successfully apply their approach to make phenotypic predictions for two E. coli strains that differ in their surface area to volume ratio. These predictions are largely consistent with various experimental datasets.

Overall, I find this work excellent. The theory provides an important new perspective and adds a valuable building block for analyzing economic principles in cell metabolism. The authors should be commended for carefully compiling a huge dataset required to (a) build/simulate the respective models and (b) to validate model predictions. While the cited paper of Zhuang et al. already

investigated some aspects of membrane crowding in combination with metabolic modeling, the framework developed here is clearly more comprehensive and advances model-driven investigations of relationships between limited membrane area and metabolic phenotypes.

RESPONSE: We thank the reviewer for the supportive comments. We truly appreciate your investment of time and your expertise. We appreciate your careful reading of the document. It keeps us motivated as we navigate the challenging world of academic publishing.

I have only few points that could be addressed in a revised version. Major points:

1) I’m a bit confused how the authors defined and treated the maintenance energy term (qATP). They defined it a as the sum of the growth (GAM) and non-growth (NGAM) associated maintenance energy demand. NGAM is clear, but what energy demands do the authors identify with GAM? On page 18 they write that ATP demand for monomer synthesis and polymerizations is already included in the biomass reaction, hence, it is not part of GAM. Please define explicitly what ATP requirements you mean by (growth-rate dependent) GAM (different from NGAM).

RESPONSE: Thank you for the comment. We define maintenance energy as an expenditure of cellular energy not directly associated with the synthesis of macromolecules (page 8). The energy required to polymerize monomers into polymers is directly associated with macromolecule synthesis and is therefore not GAM. GAM is a parameter fit between the actual measured substrate consumption and the substrate consumption required to synthesize just biomass macromolecules. NGAM is the zero growth rate intercept. We have added additional references to the document to provide readers with a suite of maintenance energy studies.

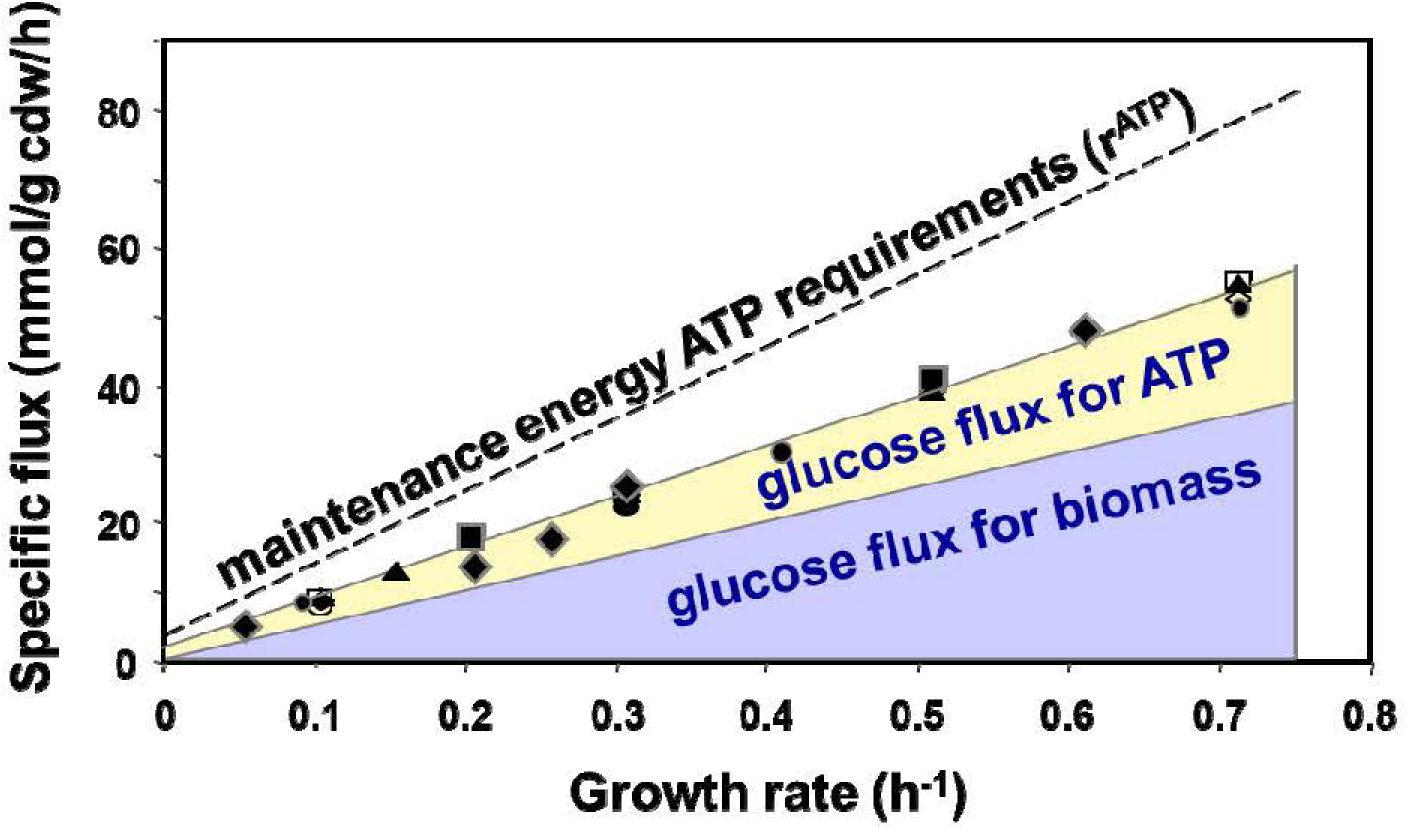

The figure above is a demonstration of maintenance energy calculation which we published previously. Symbols represent experimental glucose fluxes from: Schultz and Lipe, 1964; Neijssel et al., 1980; Tempest and Neijssel, 1987; Snoep et al., 1993; Lendenmann, 1994; de Graf et al., 1999; Sauer et al., 1999; Alexeeva et al., 2000; Abdel-Hamid et al., 2001; Emmerling et al., 2002. The experimental glucose fluxes are larger than what is needed solely for biomass synthesis (which includes monomer polymerization energy). The difference between the experimental glucose flux and the fluxes required for biomass macromolecules synthesis is assigned to “maintenance energy fluxes”. The quantitative relationship between the excess glucose consumption and ATP synthesis requires assumptions about respiration efficiency (P/O number). The intercept of the maintenance energy trend and the y-axis at a growth rate of 0 is the NGAM. Analysis from 10.1002/bit.20044.

Irrespective of the actual GAM definition, I did not understand why the authors treated qATP as independent variable. Since the authors declared that some GAM is part of qATP, there will be some ATP maintenance demand that increases with the growth rate, hence, qATP will not be independent of the growth rate. I therefore feel that Fig. 4A is not correct as it will contain regions which are not feasible (e.g. with a high growth rate and a zero or low qATP). Why aren’t the authors dealing with (N)GAM as usual: all GAM is integrated in the biomass reaction and an ATPm reaction is used to quantify NGAM. The lower bound of ATPm should be set to a typical value (e.g. 3.15 mmol/gDW/h) and larger fluxes of ATPm would indicate that additional ATP can be produced. I feel that the investigations and discussion regarding qATP/ATPm would be more consistent and some figures easier to read.

RESPONSE: Thank you for the comment. Figure 4A uses an unbiased survey of potential maintenance energy fluxes to interrogate the *in silico* role of a constrained membrane surface area on potential phenotypes. We use an unbiased survey of values because maintenance energy fluxes can change substantially between culturing conditions and can change substantially depending on how the value is defined. ‘Hardwiring’ a GAM into the biomass synthesis reaction could be problematic as the experimentally calculated ‘maintenance energy’ changes with growth conditions and with assumptions about respiration efficiency (P/O number). For example, calculated maintenance energy fluxes change by ∼ 2 fold between aerobic and anaerobic conditions in *E. coli* (10.1038/s41467-023-39724-7; 10.1016/j.cels.2016.08.013). If an aerobic GAM were hardwired into the biomass reaction, simulations examining anaerobic growth would be negatively impacted. Anaerobic fluxes and ATP yields may not be sufficient to meet aerobic maintenance energy fluxes resulting in infeasible simulations. We have added additional explanations and references for this approach in the text (p. 6).

When analyzing the experimental fluxes (Figure 5A, 6A), we recalculated GAM+NGAM for each simulation using experimental consensus data and the SA:V and protein crowding parameters. This is essential for our approach and avoids potentially problematic assumptions related to hard wiring maintenance energy values. The NGAM is the intercept of the aggregate maintenance energy fluxes when the specific growth rate is 0/h. The simulation data for MG1655 presented here has an NGAM of ∼ 4 mmol ATP (g cdw H), consistent with the reviewer’s suggested value.

2) In the manuscript, the authors used for many simulations an f_MSA of 0.07. Could you more explicitly state why you are using this value (I guess it is mentioned on page 9, but only indirectly; besides, an f_MSA of 0.07 is also used earlier on page 7).

RESPONSE: Thank you for the comment. f_MSA is the fraction of the membrane surface area occupied by central metabolism enzymes. An f_MSA = 0.07 corresponds with onset of the Pareto front between biomass synthesis and maintenance energy fluxes (Figure 5D,6C). This value was demonstrated to be accurate for both *E. coli* MG1655 and NCM3722 at interpreting the maximum observed growth rate. This makes f_MSA = 0.07 a convenient reference point.

The presented analysis can generate data for whatever f_MSA is of interest to a scientist. We have reworded this section of the manuscript to further explain the motivation (p. 6).

3) In the last paragraph of the discussion section, the authors could discuss perspectives of how to bring together their own modeling concept with other modeling approaches that explicitly include constraints for limiting resources (such as enzyme-constrained FBA models (e.g. GECKO etc.)). Wouldn’t it be straightforward to combine these different types of constraints in one COBRA models, and then to analyze how they shape together the solution space?

RESPONSE: Thank you for the comment. The presented approach is very flexible and can be integrated with other systems biology methods. The following statements with new references have been added to the discussion:

“The presented *sMSA* theory is flexible and can be readily implemented with numerous systems biology approaches including biochemical pathways analysis (e.g. EFMA) (63), resource balance analyses (64, 65), and other FBA-based approaches (42). Our research efforts have already started examining a number of these approaches.”

Minor points:

(4) Eqn. (7) in the box: how do you integrate the (growth-rate dependent) beta in the simulations given that the growth rate itself a variable in the model?

RESPONSE: Thank you for the comment. The decrease in SA:V with growth rate is assumed linear based on Figure 2B. This change in SA:V is quantitatively tracked using the beta parameter in Box 1. During a simulation, the beta parameter adjusts the available sMSA according to the growth rate. (The beta parameter is applied in a manner analogous to the classic GAM accounting mentioned by the reviewer.)

(5) Fig. 2A. The unit 1/h for overflow is a bit confusing. I understand what you want to express but perhaps you can rephrase it as: “overflow starts at growth rate [1/h]” or something similar.

RESPONSE: Thank you for the comment, we reworded this column header to ‘overflow initiation’ for clarity.

(6) I found it somewhat old-fashioned and inconvenient that the figures were not embedded in the text and that, moreover, the figure legends were places in yet another place. I think this is no longer necessary when submitting a manuscript for review.

RESPONSE: Thank you for the comment. We have incorporated the figures and captions into the main text which is now hosted in Overleaf/Latex.

(7) Shouldn’t the substantial Supplementary Material be part of the actual submission (and later be published together with the actual paper), instead of being hosted in a github repository?

RESPONSE: Thank you for the comment. We would be happy to upload the supplementary materials when the manuscript finds a home for publication. In the meanwhile, we are storing it on github.

XXXXXXXXXXXXXXXXXXXXXXXXXXXXXXXXXXXXXXXXXXXXXXXXXXXXX XXXXXXXXXXXX

Cell Systems

Manuscript submitted: 6 June 2024

Reviews received: 9 December 2024

Cell Systems Reviewers’ comments:

Reviewer #1: Review of the paper by Ross P. Carlson et al. Summary of the work:

In this work authors presented a ‘metabolic theory’, based on microbial cell geometry, membrane protein crowding and metabolism to predict growth rate, overflow metabolism and maintenance energy flux. To emphasize the contribution from cell geometry, authors claimed the capacity of cell membrane to host proteins increases with growth rate and compensates for the decrease in surface area to volume ratio with increasing growth rate. To support their membrane centric hypothesis, authors used NCM3722 and MG1655 strains, which authors claimed to be genetically similar but differ in surface to volume ratio, maximum growth rate and overflow metabolism phenotypes. In this study authors did not consider any contribution of cytoplasmic crowding to affect the maximal growth rate or other phenotypes. In this work, authors concluded that cell-geometry and membrane protein crowding as significant regulator of cellular physiology.

The authors tried to build a membrane area centric predictive model to explain cellular behaviors related to maximal growth rate and overflow metabolism.

However, the work fails to demonstrate predictive power of the model. The experimental perturbations that are used to validate the model are insufficient to test model predictions and are too similar to data used for a basic model parameter calibration.

Even more concerning, the major conclusions of this model are in direct contradiction with a long literature about enhancing growth rates and perturbations that affect S/V, but do not affect growth rates.

Unfortunately, we do not find this work convincing overall we do not think this work is suitable for publication in Cell Systems. These major issues that cannot be easily overcome by revising the manuscript.

RESPONSE: We would like to thank the reviewer for their investment of time reading and commenting on the manuscript. We appreciate the opportunity to sharpen our arguments, identify additional data that support our hypotheses, and refine the presentation of the results. We have removed ∼25% of the manuscript figures to focus on the major elements of the theory. We have identified additional experimental data that directly support our hypothesis; changing the SA:V ratio of *E. coli* MG1655 using titratable MreB expression changes the growth rate in a predictable manner consistent with the presented theory. We have also reorganized the results section so that MG1655 data is used to identify five predictions which are then tested using independent NCM3722 data. These predictions are as follows: **prediction 1** *sMSA-FBA* using an *f_MSA_* of 0.07 will predict *μ_max_* for NCM3722 at the Pareto front between biomass flux and maintenance energy flux, **prediction 2** *sMSA-FBA* will only predict an accurate *μ_max_* for NCM3722 when biologically relevant SA:V ratios are applied to the analyses, **prediction 3** *sMSA-FBA* will quantitatively predict NCM3722 P/O ratios highlighting the optimal use of finite membrane surface area and protein crowding, **prediction 4** *in silico* NCM3722 P/O ratios, plotted as a function of growth rate, will have an inflection point that corresponds with the experimental onset of acetate overflow, and **prediction 5** the most efficient NCM3722 biomass yield on glucose will correspond with the inflection point of P/O number vs. growth rate providing a basis for defining an optimal glucose phenotype. The five predictions are demonstrated to be accurate using data from independent studies and a different *E. coli* strain (Figure 6). We believe the presented data and arguments support our theory that cell membrane processes are major contributors to phenotype.

Major concerns:

- Genetically, controlling expression levels of FtsZ changes cell length several-fold. Surface to volume ratio changes much more under this perturbation than for the conditions considered in this manuscript. However, growth rate is unchanged across the entire range of FtsZ titration. This experimental result is in direct contradiction with the model proposed in this work. Similarly, there are cell wall mutants that affect cell width and have a strong effect on surface to volume ratio and have undetectable effects on growth rate.

RESPONSE: Thank you for the comments and pointing us in this interesting direction. We have identified an experimental data set that directly supports our theory and hypotheses. Si et al built an MG1655 strain with a titratable MreB.

MreB alters the cell diameter and thus the cell SA:V. The truly remarkable data set demonstrates a positive relationship between the engineered strain’s SA:V and its growth rate on defined medium. The data is consistent with the presented theory and provides a direct test of the hypotheses in this work. MreB is generally believed to influence cell diameter while FtsZ influences cell length. The curvature of the cell is likely a key element of protein hosting which is positively influenced by MreB but not FtsZ. Interestingly, when this data was plotted as the total cell volume vs. growth rate, there was a negative trend. Thank you again for the suggestion, much appreciated!

- It has also been shown previously, that in E. Coli growth rate improvements can be achieved on many substrates. For example, point mutation in glycerol kinase glpK, results in growth rates of E. Coli on glycerol comparable to those glucose. Similarly, Cra knockout and overexpression of ArcA result in faster growth rates on many substrates. These experimental data suggest that growth rates are not maximized in most conditions.

RESPONSE: Thank you for the comment. These experimental observations are consistent with the predicted framework. As noted in the manuscript, *E. coli* does not operate at its maximum theoretical growth rate; it grows at a rate substantially lower than what is theoretically possible due to maintenance energy demands (Figure 3). We present the physiological limits of *E. coli* based on its central metabolism, SA:V ratio, and membrane protein crowding. Computational biology dogma of assuming a ‘maximum growth rate’ is a misnomer. The maximum growth rate is, in actuality, a relative maximum growth rate due to using a maintenance energy drain to fit experimental data. We state this in the document and quantify the slope of the maintenance energy vs growth rate relationship while holding biomass yield constant. Gene deletions or evolved strains that reduce maintenance energy demands/fluxes (such as removing potential futile cycling e.g. glycerol and glycerol-P, futile cycling involved with sensing nutrient levels glutamate/glutamine, etc) could increase growth rate. Of course, these specialized mutant strains will likely sacrifice wild type robustness for the benefit of faster growth rates under certain culturing conditions.

- Even without the points above, there are real experimental tests of the model. Most of the data presented is from different carbon limited growth rates. It’s not so surprising that a model tailored to these conditions can match a few data

points. However, none of these plots constitute a serious test of model predictions. Thus predictive power of the model has not been demonstrated.

RESPONSE: Thank you for the thoughtful comments. We have reorganized the results section. The wildtype MG1655 data is now presented first and is used to define the theory and to develop five hypotheses. The latter half of the results then tests the five hypotheses using independent *E. coli* NCM3722 data. The predictions are remarkably accurate as shown in Figure 6.

- “We demonstrate that the capacity of the membrane to host proteins increases with growth rate offsetting decreases in surface area-to-volume ratios (SA:V).” There is a simple interpretation, which is that membrane capacity is not limiting in any of these conditions.

RESPONSE: Thank you for the comment and challenge. We have moved important data from the supplementary data to the main document to highlight the dynamic nature of the membrane proteome. This data was critical to our development of the presented theory. Evolution selects competitive phenotypes. Dynamic changes in the membrane proteome would not be necessary if the membrane proteome had no impact on fitness. If the membrane protein capacity was not limiting, we would predict the membrane catalytic capacity would be invariant with growth rate or would even decrease as those resources could be directed to cytosolic components like ribosomes. Secondly, we present additional experimental data from Si et al. demonstrating that an increase in SA:V increases the MG1655 growth rate. Additionally, the discussion highlights how bacterial cells, which have evolved to become highly specialized in respiration and ATP generation, have also evolved to host a larger protein content in their membranes. These highly specialized, highly evolved bacteria are also known as mitochondria.

- The presented theory doesn’t take cytosolic crowding into account. However, it has been shown that overexpression of purely cytosolic, non-membrane proteins shifts the acetate excretion threshold.

RESPONSE: Thank for the comment and the opportunity to discuss this work. The referenced study surely contributes to the field’s knowledge and certainly modulated cytosolic crowding, but it is important to recall the data was collected from a recombinant strain. Carlson would argue these modifications open the door to other possible interpretations of the data. Carlson would like to respectfully highlight aspects of that study which may permit additional interpretations. The referenced strain’s modifications necessarily perturb the very phenotype that was the subject of study. 1. The strain perturbed native gene expression by removing a gene central to native glucose metabolism and therefore central metabolism. 2. The strain perturbed the native membrane proteome by recombinantly expressing only one subunit from a large multisubunit enzyme complex (PtsG, Crr, PtsI, PtsH). 3. The strain perturbed the native metabolic regulation of glucose metabolism by disconnecting cell growth rate and cellular energy state (ATP/ADP) from the sugar transporter saturation (S/(S + Km)). The referenced study controlled growth rate via recombinant expression of a protein, yet the extracellular glucose concentrations were many orders of magnitude higher than the transporter Km value. Glucose is a preferred *E. coli* substrate and the Pts system is both a transporter and a sensor that regulates many metabolic functions (e.g. cAMP/CRP, ArcA, etc regulated processes). 4.

The cytosolic proteome was perturbed through the recombinant expression of an additional protein. 5. The macromolecular composition of the cell was perturbed through the expression of multiple recombinant proteins. 6. The strain maintained a recombinant plasmid(s) and expressed antibiotic resistance protein(s). 7. The cellular carbon, nitrogen, and reducing equivalent pools were all perturbed through the combination of issues 1-6. These perturbations would potentially influence a wide range of other regulatory and metabolic features. However, the current study was able to reconstruct many of the salient phenotypic features of the recombinant strain using only the presented sMSA criteria. Carlson does not see the interpretation of the referenced study and presented study as a zero sum game. Ultimately, a cell needs to balance both membrane and cytosolic processes to maximize fitness. The two studies are complementary.

Minor comments:

In figure 2, panel 2B and 2C are not matching with the text presented in figure legend.

RESPONSE: Thank you for the careful reading of the document. The captions have been corrected.

Reviewer #2: This paper presents a theory about membrane occupancy of proteins important for central metabolism and explores its implications through FBA simulations that incorporate a constraint formulated based on the theory. The topic that is addressed by this paper has gained considerable interests for more than a decade now. However, I find it difficult to identify the contributions of this work compared to the previous research.

RESPONSE: Thank you for reading and commenting on our manuscript. Your efforts are much appreciated.

My main concerns/comments are:

(1) While the simulation analysis of this work has led to propositions regarding the potential role of the membrane surface area constraint in shaping the growth rate, biomass yield and overflow (including the impact of maintenance energy), it is unclear how the positions stated in this work are fundamentally different from those given by previous work on membrane surface limits (e.g. Zhuang et al 2011, Szenk et al., 2017).

REPSONSE: Thank you for carefully reading the document and the comment. We have shortened and rearranged elements of the introduction to emphasize and clarify the advances presented here. The introduction makes the following statement to support the novelty of the results. “While theoretical studies have addressed the role of surface area on metabolism (15, 16), no study has developed a quantitative and predictive molecular level theory that accounts for strain-specific differences in SA:V ratios, growth rate dependent changes in SA:V ratios, and growth rate dependent changes in membrane protein crowding (29). The presented systems biology theory demonstrates the remarkable theoretical impact of cell geometry and membrane protein crowding on foundational cellular processes including maximum growth rate, onset of overflow metabolism, respiration efficiency, and ultimately the cellular capacity for maintenance energy generation. These predictions are all accomplished without consideration of cytosolic macromolecular crowding nor absolute cellular volume highlighting gaps in current systems biology theory.” These aspects are again highlighted in the discussion section to reinforce the novelty and transformative potential of the study.

(2) A key point made by the paper seems to be on (a) the decrease of SA:V ratio as growth rate increases and (b) the compensation for this decrease by the increase in the capacity of the membrane to host proteins. I wonder however how these two factors are represented in the specific membrane surface area constraint (eqn. 6 in Box 1): are they combined into the parameter beta? And more importantly, how different is this constraint from that used in Zhuang et al. in terms of the actual impact on FBA simulation results?

RESPONSE: Thank you for the comment and the careful reading of Box 1. The SA:V ratio is introduced in equation 2 as the experimentally measured equation of form c1*mu + c2 (from Figure 2B). The protein crowding, quantified as fMSA, is introduced in equation 3. Zhuang et al do not account for growth rate dependent changes in the SA:V nor does it account for the change in protein crowing with growth rate. These parameters are demonstrated to be essential to the presented results. We have reworded portions of Box 1 and the theory development section of the manuscript to clarify the importance and implementation of these central parameters.

(3) In Discussion, the authors hypothesize that “both surface area- and volume-associated processes are critical and concurrently influence phenotype”. I wonder whether this work has shed new light on what phenotypical aspects (if any) are primarily governed by the surface area-based (as opposed to other) constraint. In the simulation results obtained in this work, did the surface area-based constraint represent “the” governing mechanism, or rather one of the possible mechanisms? Does this work provide insights on how the limits posed by the available surface area relate to those by volume or by the capacity of the protein expression machinery in the governing of phenotype?

RESPONSE: Thank you for the comment and providing the opportunity to strengthen our work. This work develops a new dimension to our understanding of cell biology and provides a basis to answer basic questions that have confounded scientists for decades. For example, the P/O number is quantitatively predicted here, found to accurately align with experimental values, and the theory demonstrates why these values, which have been reported for decades, represent a competitive phenotype. These predictions and interpretations would not be possible from only examining the cytosolic crowding as the enzymes responsible for the P/O number are embedded in the membrane. The current study also provides a biophysical basis for calculating maintenance energy using constraints associated with the membrane. Maintenance energy accounts for 30-90+% of cellular fluxes. These substantial drains on metabolism are typically handled in computational models by simply using a generic fitting parameter. That is no longer necessary if the membrane SA:V and fMSA parameters are defined.

Minor comments:

(4) Title: “Cell geometry” is mentioned there; I am not sure how important it has been in this work.

RESPONSE: Thank you for the comment. We have added the descriptor “cell geometry, as quantified by cell shape and cell dimensions” to highlight the essential aspects of both cell shape and dimensions to the presented work. The geometry based surface area and cell volume equations are functions of both the cell shape and dimensions.

(5) Page 5, bottom: “We…developed a…theory to test the hypothesis”: Could one test a hypothesis using a theory?

RESPONSE: Thank you for the comment. We have reworded the sentence and reorganized the results section into a theory development section and a theory testing section.

(6) Page 6, end of 2nd paragraph: Some past studies with the membrane constraint (e.g. Ref [13], Zhuang et al.) were already expressed in terms of area, not mass.

RESPONSE: Thank you for the comment. You are correct; however, the Zhuang et al study used qualitative, representative values for the enzyme properties - as the detailed values used here had not yet been measured. The current study takes advantage of the new data generated during the last decade+ to create a quantitative and predictive computational approach. We state the following in the introduction: “While theoretical studies have addressed the role of surface area on metabolism (15, 16), no study has developed a quantitative and predictive molecular level theory that accounts for strain-specific differences in SA:V ratios,

growth rate dependent changes in SA:V ratios, and growth rate dependent changes in membrane protein crowding (29). The presented systems biology theory demonstrates the remarkable theoretical impact of cell geometry and membrane protein crowding on foundational cellular processes including maximum growth rate, onset of overflow metabolism, respiration efficiency, and ultimately the cellular capacity for maintenance energy generation. These predictions are all accomplished without consideration of cytosolic macromolecular crowding nor absolute cellular volume highlighting gaps in current systems biology theory.”

(7) Page 6, close to bottom: “ETC”: Electron Transfer Chain?

REPSONSE: Thank you for the careful reading of the document. We have added a definition of ETC.

(8) Page 9, close to bottom, the subsection title says “Maximum growth rate maximizes the areal density of ATP synthase and rate of ATP hydrolysis”: I wonder whether the specific ATP cost for growth was kept constant in all the simulations. If so, wouldn’t this statement be rather obvious?

RESPONSE: Thank you for the comment. We have removed this analysis to avoid distractions and now focus on fewer topics but in more detail. However, to answer the question, no, the specific ATP cost was not fixed in these simulations. As noted, to do so would be obvious.

(9) Page 10, middle, subsection title, “energy conserving potential”: what “conserving potential” means here?

RESPONSE: Thank you for the comment and opportunity to clarify our work. A sentence has been added that states “These enzymes can conserve varying amounts of substrate chemical energy by converting it into chemiosmotic energy stored as proton motive force.”

(10) Page 26, Fig 2, “P/O” should be marked on plot B and “PtsG” on plot C?

RESPONSE: Thank you for the careful reading of the document. We appreciate your attention to detail. We have corrected the figure references.

XXXXXXXXXXXXXXXXXXXXXXXXXXXXXXXXXXXXXXXXXXXXXXXXXXXXX XXXXXXXXXXXX

PNAS

Manuscript submitted: 8 March 2024

Reviews received: 14 April 2024 Reviewer Comments:

Reviewer #1:

Suitable Quality?: Yes

Sufficient General Interest?: Yes Conclusions Justified?: Yes Clearly Written?: Yes Procedures Described?: No

Comments:

Beg et al published a paper in PNAS (PMID: 17652176) where this paper demonstrated that molecular crowding is an important constraint in E. coli metabolism. The concept of molecular crowding is adopted in ME-model for E. coli at MSB (PMID: 24084808), where this paper demonstrated the concept of proteome limitation constraint can be hit at the maximum growth rate and can explain the metabolic shift in E. coli. Furthermore, Basen et explained the metabolic shift in E. Coli can be explained based on the proteome allocation and published their study in Nature (PMID: 26632588). After that Vazquez and Oltvai used Basen’s model and they evaluated their molecular crowding constraint at PNAS, and they reported how the molecular crowing constraint can also explain the metabolic shift (PMID: 27484619). In conclusion, these studies showed that the molecular crowding constraint is important in E. coli.

On the other hand, Zhuang et al showed that the cellular membrane can be an important constraint that explains the metabolic shift in E. coli and published their study at MSB (PMID: 21694717), where the glucose transporter and cytochrome oxidase and the other OxPhos complexes can have competition because the available area of membrane is limited. Furthermore, Szenk et al wrote a perspective at Cell Systems (PMID: 28755958) that supported the limitation in the cellular membrane because of surface-to-volume ratio decreases at higher growth rates.

ME-model combines these constraints by predicting the membrane constraint is hit at the low growth rates and the proteome limitation is hit at the higher growth rates.

In this manuscript, Carlson et al collected biophysical data and proteomics data and combined them with metabolic fluxes to show the importance of surface-to-volume constraint in three E. coli strains. Their predictions support Szenk et al’s perspective, however, there are some issues in the authors’ metabolic model.

The authors should compare the predicted O2 and CO2 fluxes with the measured fluxes. This comparison is important to reveal the GAM and NGAM values are used correctly in the model. Additionally, the authors should explain the biological meaning of the constrain fMSA = 0.07 and the biological meaning of the parameter 0.07. Please see below my comments in detail:

RESPONSE: Thank you for the nice summary of relevant literature. We have used this information to edit our introduction and to cast the membrane protein crowding as a major feature of the analysis.

The comments about CO2/O2 and the fMSA are addressed below, briefly, the presented model is atom and electron balanced (supplementary material S5). When the biomass synthesis rate and acetate secretion rate are accurate, it is mathematically necessary that the CO2 and O2 fluxes be accurate. The CO2 is required to balance the carbon and the O2 is required to balance the electrons. The biological interpretation of fMSA is discussed below when the comment is raised there.

1- I did not read Supplementary Text S1, because this file is not on GitHub, therefore I could not review this important file.

RESPONSE: Thank you for the careful reading of the document. We have moved this material from supplementary material to the main document as Box 1 to ensure all theoretical material is available to a reader.

2- The main goal of the manuscript is to define the “specific membrane surface area (sMSA) as the area occupied by membranal enzymes per gDW (nm^2/gDW). The authors applied a sensitive analysis to estimate the total fraction of this occupied area per surface area (fMSA) between 0.04 to 0.08 (Figure 3). The biological meaning of fMSA is not clear in the text.

RESPONSE: Thank you for the comment. We have reworded the section of text and the material in Box 1 to highlight the definition of fMSA as the fraction of the surface area occupied by central metabolism enzymes. The original Figure 3 has been removed to streamline the presentation of the material. Figures 5 and 6 now present the case for an fMSA of 0.07 in a more concise manner.

How did the authors convert gdw to cell volume? The authors then predicted the surface area from the cell volume. The role of the molecular crowding constraint is included in this step. As I missed the supplementary text 1, I think that the authors used the cell density parameter to convert cell mass to cell volume.

Otherwise, the authors convert the growth rate to the volume and surface area based on Figure 1. In both cases, the role of molecular crowding constraint can play a key role in the author’s calculations. However, the authors did not mention anything about the molecular crowding constraint in the text. The use of the cell density parameter means that the authors assume that the molecular constraint is active.

Therefore, the authors should clearly explain the biological meaning of their parameter value fMSA = 0.07 in Figures 3 and 4.

RESPONSE: Thank you for carefully reading of our document. The conversion between cell volume and g cdw is developed in Box 1. Cell volumes can be converted to g cdw using the density of a live wet cell and the percentage of a cell which is water. The conversions are shown in Box 1 while a survey of literature values is presented in the supplementary material S4. The model accounts for the change in cell size with growth rate to ensure accurate conversions with growth rate.

3- The second important parameter in the text is the fraction of proteome per lipid membrane ratio. Based on the authors’ supplementary data S4, they used the median value as the proteome fraction can cover 24% of the surface area. This fraction is close to mammalian cells that grow at lower growth rates than E. Coli.

The authors reported also that the mitochondrial proteome membrane can cover 44% of the surface area. Furthermore, in another missing reference, the outer mitochondrial proteome membrane can cover 50% of the surface area, while the inner mitochondrial proteome constraint can cover 80% of the surface area of the inner mitochondrial membrane (https://web.archive.org/web/20151123055452id_/ http://www.med.ufro.cl:80/clases_apuntes/cs_preclinicas/mg-fisica-medica/sub-modulo-1/Mitochondria.pdf)

RESPONSE: Thank you for the information. We have updated the discussion and added data to the parameter survey in supplementary material S4. Please note the referenced article and many other eukaryotic studies do not use ratios of areas but instead ratios of molecules.

Additionally, Zhuang et al (PMID: 21694717), used the fraction of proteome to cover about 50% of the surface area. From these data, the proteome can cover about 50% of the surface area which is close to the reported value by the authors for the mitochondria.

Therefore, the authors should carefully identify the value of this important parameter.

RESPONSE: Thank you for the comment. We present a survey of published membrane surface area values in supplementary data S4. pubs.acs.org/doi/full/10.1021/jp8107446, www.nature.com/articles/ncb2561, doi.org/10.1016/j.cell.2006.10.030, www.pnas.org/doi/abs/10.1073/pnas.0712379105, doi.org/10.1038/s41598-017-16865-6, pubs.acs.org/doi/full/10.1021/ja902853g, pubs.acs.org/doi/full/10.1021/jp8107446, doi.org/10.1016/S0005-2736(00)00323-0

The median value for the literature review is 24+/-5.6% of the membrane area is occupied by protein. The value in Zhuang was a proposed value based on mitochondria and was published before more recent E. coli proteomics studies were conducted. Mitochondria have the luxury of existing in a buffered cytosol and can support higher protein levels. Free living bacteria typically persist in harsher environments requiring different membrane strategies.

4- Page 6, the authors wrote that “fMSA = 0.04-0.08 (4-8% of surface area occupied by central metabolism enzymes, supplementary text S1)”.

By dividing fMSA by 0.24 (proteome fraction in surface area, the value that the authors used), therefore the model proteome can cover only 33% of the available proteome in the surface area. If they used 50% of the surface area occupied by proteins (as mentioned in the previous comment), fMSA represents only about 17% of the available area of the proteins.

Based on these calculations, the cell can increase the proteins in the surface area because of the available area for proteins. Therefore, a reader can understand that the surface area constraint is not active in these conditions. The low value of fMSA indicates that there is another active constraint that prevents increasing fMSA value. However, the authors did not mention this constraint that prevents fMSA from increasing.

Therefore, based on the previous comments the authors should explain their constraint (fMSA =0.07). They also should discuss the biological meaning of their constraint and the parameter values.

RESPONSE: Thank you for this interesting question. The fraction of the surface area occupied by central metabolism enzymes is less than the total available surface area. This is based on numerous studies, for instance: Belliveau NM, et al. (2021) Fundamental limits on the rate of bacterial growth and their influence on proteomic composition. Cell Syst 12(9):924-944 e922. Schmidt A, et al. (2016) The quantitative and condition-dependent Escherichia coli proteome. Nat Biotechnol 34(1):104-110. Valgepea K, Adamberg K, Seiman A, & Vilu R (2013) Escherichia coli achieves faster growth by increasing catalytic and translation rates of proteins. Mol Biosyst 9(9):2344-2358. Peebo K, et al. (2015) Proteome reallocation in Escherichia coli with increasing specific growth rate. Mol Biosyst 11(4):1184-1193. Papanastasiou M, et al. (2013) The Escherichia coli peripheral inner membrane proteome. Mol Cell Proteomics 12(3):599-610.

We hypothesize there are essential structural proteins like MreB and FtsZ and additional transporter proteins like aquaporins, quorum sensing transporters, ion channels, etc. that occupy the rest of the area precluding all of it from being used for central metabolism. These requirements can be added to the presented theory as the data becomes available

5- The authors used the biomass reaction in their model as “0.95 acetyl_CoA + 36.9 ATP + 0.41 D-erythrose_4-phosphate + 0.05 a-D-glucose_6-phosphate + 4.4 L-glutamate + 2.77 L-glutamine + 26.83 H2O + 3.26 NAD + 13.53 NADPH + 1.88 oxaloacetate + 2.1 phosphoenolpyruvate + 3.13 pyruvate + 0.69 D-ribose_5-phosphate + 0.5 S2H + 6.28 MRE ==> 36.9 ADP + 6.04 2-oxoglutarate + biomass(1kgcdw) + 0.95 coenzyme_A + 22.88 H+_cyt + 3.26 NADH + 13.53 NADP + 38.93 phosphate”

There are some questions about this biomass reaction. First, the GAM is 36.9. This value is lower than the GAM value used in the E. coli metabolic models which is close to 60 mmol/gDW (check Table 15 from Palsson’s group in https://journals.asm.org/doi/full/10.1128/ecosalplus.10.2.1). The lower GAM value in the model leads to that the model did not demand more membrane area for oxygen in respiration or glucose in the fermentation phases.

Secondly, the biomass compositions differ from Table 15 in the previous reference for core metabolism in E. coli.

Finally, the authors added the MRE as a substrate in the biomass reaction. The unit of biomass compositions in the biomass reaction must be mmol/gDW. What is the unit of MRE? It is area per gDW. In this case, the definition of the add operation is not satisfied (by adding two distinct types). Therefore, the authors should adopt the formulation from the GECKO model from Nielsen’s lab at MSB (PMID: 28779005) by assuming that there is amount of enzyme (mmol/gDW) is shared in a specific reaction. The authors then can define the membrane constraint as the protein mass constraint in the GECKO model.

RESPONSE: Thank you for the careful analysis of the metabolic model and biomass reaction. The biomass reaction does not include the growth associated maintenance energy (GAM) nor does it include NGAM. The biomass reaction only includes the ATP required to polymerize monomers into macromolecules.

We have reworded the model description to highlight this important detail.

Secondly, the biomass stoichiometry is based on our previous work and the work of the Uwe Sauer group, as referenced in the document. The biomass stoichiometry has been analyzed for electron density (degree of reduction) and is consistent with experimental biomass measurements. We have published analyses on biomass reactions and are confident it is a reasonable representation of the metabolic demands to produce biomass. 10.3390/pr6050038

Finally, the MRE balance is orthogonal to the mass balance on metabolites. It is based on the surface area available per g cdw. We have reworded the model section to highlight the units and the orthogonal nature of the balance.

6- The authors plotted the ATP synthase areal density (complexes per micro m^2) in Figures 3 and 4. However, the authors did not mention the glucose, oxygen, and acetate fluxes in the main text to validate that their protein abundances can satisfy the experimental flux values, especially the oxygen fluxes. The comparison between oxygen and CO2 fluxes between the model predictions and the measured fluxes is especially important to reveal the role of GAM and NGAM in the model.

RESPONSE: Thank you for the comment. The model reactions are all balanced for atoms and for electrons. The model only has oxygen as an external electron acceptor while overflow and fermentation products can also serve as sink electrons. The balanced reactions are shown in supplementary material S5. The predictions are therefore carbon and electron balanced so with the simulations having accurate biomass and acetate fluxes, the CO2 production rate (satisfies carbon balance) and oxygen consumption rate (satisfies electron balance) are consistent with experimental data since the model has no other means of balancing these compounds.

7- The authors define some reactions in the TCA pathway, ENO reaction, and carbon transporters as D-glucose_ext + phosphoenolpyruvate + x MRE + 6 Cmol + 24 Emol + Mol ==> a-D-glucose_6-phosphate + pyruvate acetate_ext + H+_ext + x MRE + 2 Cmol + 8 Emol + Mol ==> acetate + H+_cyt Cmol_ext ==> Cmol

The authors add the metabolite Cmol to these reactions. To the best of my knowledge, I did not see this formulation in metabolic models. The authors couple the glucose, acetate, and TCA reactions by the flux Cmol_ext ==> Cmol. In this case, the authors assume a specific constraint that can couple between these fluxes. What is the biological meaning of this constraint? The reason for this formulation is not clear in the text.

RESPONSE: Thank you once again for the careful analysis of the supplementary material. We appreciate working with a reviewer who understands the importance of all the parameters and modeling elements. The MRE balance accounts for the quantity of surface area available per g cdw, as defined in Box 1. The model includes terms that permit the tracking and quantification of electron moles (Emol), carbon moles (Cmol), and total metabolite (Mol) moving across the cell membrane. These theoretical metabolites are not used as constraints for the analyses but are used to examine trends related to membrane trafficking of different resources. We have updated supplementary data S5 to denote the identity of the model feature and to denote that these theoretical metabolites are not used to constrain results.

8- There is a problem with Figures 1A and 1D. First, the growth rates in Figure 1A are between zero and 1.5 per hour, but the growth rates in Figure 1D are between zero and 0.58 per hour. As the authors wrote “While the SA:V ratio decreases with growth rate, the capacity of the membrane to host proteins per cell volume increases with growth rate (Fig. 1D).” Figure 1D lacks what can happen in the membrane proteome between the growth rates between 0.58 and 1.5 per hour, although Schmid et al paper reported proteomics data until the growth rate 1.5 per hour (PMID: 26641532 Figure 2). Therefore, the authors’ hypothesis may be limited until the growth rate of 0.58 because we do not know the data between the growth rates of 0.58 and 1.5 per hour.

RESPONSE: Thank you for the comment. The data was collected under different growth conditions. Some data is from glucose minimal medium which limits the mu max to ∼0.7/h while cultures grown on complex or semi-complex medium grew much faster (∼1.5/h). This explanation is highlighted in the document.

9- In Figure 1D the authors normalized the area occupied by proteome with surface area, why did not authors normalize this area occupied by proteome with surface area? Therefore, the reader can understand Figure 1E with different growth rates. Furthermore, figure 1E has proteomics data for the growth rate of 0.7 per hour, whereas this data is not available in Figure 1D.

RESPONSE: Thank you for the comment. We have replotted the data so that it is all on a per volume basis. This was selected so we can compare cell SA:V trends (Figure 2B) with protein occupied area per volume (Figure 3A) to appreciate the role of enhanced membrane protein crowding as a function of growth rate. The text now mentions this basis for normalizing the data.

10- Page 6, the authors wrote “Note, the lower right portion of Fig. 2F is a region of extreme sMSA scarcity with predicted lactate overflow.” The authors predicted acetate and did not predict lactate.

RESPONSE: Thank you for the comment. We have removed the statement to avoid introducing confusion. This section of the plot is theoretical and not observed under typical batch growth although the prediction of a minimal metabolism at the maximum possible growth rate is interesting.

11- The figures contain error bars generated from the simulations. What are the reasons for the change in the prediction values at different runs?

RESPONSE: Thank you for the comment and opportunity to improve the document. Each set of experimental consensus data was analyzed 100 times using FBA, with each simulation using a perturbed set of strain specific consensus fluxes (mean data +/- up to 1 standard deviation). The consensus fluxes are plotted in Figure 5A and 6A. The error bars from the simulations therefore represent the sensitivity of the results to perturbations of the consensus fluxes. We have reworked the model development and analysis section to highlight this feature of the FBA analysis. Additionally, the document has all used MATLAB code included in the supplementary material.

12- Authors applied curve fitting with high-order polynomials, what is the biological meaning of these polynomials?

RESPONSE: Thank you for the comment. This fitting was empirical and based on an analysis of the MG1655 and NCM3722 data. The analysis of the data can be found for both strains in supplementary material S7. Additionally, Figure 6 plots several additional P/O number curves (perturbed based on different SA:V ratios) which reinforce the nonlinear nature of the relationship.

Reviewer #2: Suitable Quality?: No

Sufficient General Interest?: No

Conclusions Justified?: No

Clearly Written?: No

Procedures Described?: No

Comments:

The manuscript describes a model of cellular metabolism and membrane protein allocation, which the goal of exploring possible relations of various cellular phenotypes, including overflow metabolism and growth, to cell size. The model is developed and validated using available flux and proteomics data for different E. coli strains characterized by different surface are to volume ratios (SA:V).

Overall, I am confused over what are the actual predictions of the model. The abstract states that “the theory successfully predicts the phenotypes of two E. coli K-12 strains, MG1655 and NCM3722, which are genetically similar but have different SA:V ratios, different maximum growth rates, and different overflow phenotypes”. This is a remarkably vague sentence, and it should be made clear -- both here and in the flow of the manuscript -- which of these features are used as model inputs and which ones are genuine predictions.

From the modelling point of view, the theory proposed by the authors appears to be quite similar to what was developed in Ref. 14, Zhuang et al., in which the effect of a constraint on membrane occupancy on FBA solutions was analyzed. I cannot access Supplementary Text S1 at the provided github link; without looking in detail at the derivation of the model, it is hard to evaluate how different the approach proposed here is from the one in Ref. 14. My understanding is that, while in Ref. 14 the constraint is on the fraction of membrane area occupied by the enzymes, here the overall membrane surface is also taken to be condition-and strain-dependent. While this is unlikely to produce qualitative changes in the predicted FBA solutions, it will lead to different quantitative predictions, which are thus crucial for evaluating the relevance of the model developed here. A crucial aspect of the model is the claimed ability of explaining the different phenotypes of E. coli NCM and MG strains. These predictions should be explained better and quantitatively supported by the data.

The flow of the manuscript could benefit from streamlining the presentation of the simulation results; it does not help that different simulations are performed using different sets of constraints and/or objective functions.

RESPONSE: Thank you for investing time into our manuscript and providing valuable insight for improving our document. We have added numerical values to the abstract statement to support the statements. We have streamlined and reorganized the results section to clarify the theory. We now have 1) a theory development section using MG1655 data, 2) a predictions section based on MG1655 data, and 3) a theory testing section using independent data for strain NCM3722. The theory does a remarkable job predicting the phenotype of NCM3722.

The current work is a major development from earlier work by Zhuang et al. This has been highlighted in the reorganized introduction. The current study explicitly integrates growth rate dependent changes in SA:V, growth rate dependent changes in the surface area available for proteins (fMSA), a quantitative treatment of the enzyme kinetics, a quantitative treatment of the surface area required per enzyme, and the quantitative comparison of two related but phenotypically different E. coli strains. These features are major advances to the presented theory based on remarkable experimental data that was not available when Zhuang was published in 2011.

The mathematical development of the theory has been moved from the supplementary material and is now in the main document as Box 1.

We have streamlined the document, removing side stories to focus on the major elements of the theory. As mentioned above, we have reorganized the results section to highlight what the predictions are and how the experimental data confirm the predictions.

Here below are some major, specific points to be addressed.

Fig. 1D - Is the areal density of membrane proteins for NCM3722 the same as for MG1655?

RESPONSE: The areal density of NCM3722 and MG1655 proteins are demonstrated to have similar values based on the presented analyses (Figure 5D and 6C). This assessment is the result of ∼12 independent published studies processed through the presented SA:V and membrane protein crowding theory. Both strains demonstrate a Pareto trade off in the capacity to produce biomass and maintenance energy when fMSA = ∼0.07. These conclusions use the same metabolic model with the same enzyme parameters but differ only in the applied SA:V and experimental consensus fluxes. The results section has been reorganized to emphasize the major conclusions of the study.

Fig. 3A - The best fit in Fig. 3A for NCM appears to be at a fraction around 4-5%, and at a somewhat larger fraction for MG1655. It would be better to show the best fit for NCM and for MG strains separately, along with plots of the fitted glucose fluxes (or growth yields) and acetate fluxes used to compute the euclidean distance. How different the predictions would be using these two best fit values?

RESPONSE: Thank you for the comment. We have removed this figure to avoid confusion. We now focus on a streamlined set of data and analyses (e.g. Figure 5 and 6). However, the analysis demonstrated that both the MG1655 and NCM3722 used similar percentages of the available surface area for central metabolism enzymes.

Fig. 3D - In this figure results are shown for simulations with a fMSA value of 0.07 common to both cell types. As MG cells are smaller, they have higher SA:V ratio and thus can fit more membrane enzymes compared to NCM3722. The ATP concentration is predicted to be higher in MG compared to NCM (Fig. 3D). So, why wouldn’t grow faster than the NCM strain and, instead, just has a higher density of ATP synthase?

RESPONSE: Thank you for the comment. We have removed this figure to streamline the presentation of the theory and to avoid confusion. However, the predictions demonstrate that the maximum growth rate occurs when the available membrane surface area and protein crowding prevent use of additional membrane associated central metabolism The NCM3722 strain is in fact smaller than MG1655 and has a higher SA:V ratio which is predicted to enable the experimentally observed faster growth rates on glucose minimal media. We have also added published experimental data that demonstrates changing the SA:V changes the mu max in a predictable manner (Figure 5F).

Fig. 3E - Of the data presented, it looks like only the dataset from Peebo et al. shows a clear reduction of ATP synthase abundance; other datasets show a range of behaviors. Are there differences between MG and BW strains? What about the NCM strain, is it indeed lower than in the MG/BW strains? This plot is crucial to support the hypothesis of the maximal growth being associated to the maximal allocation of ATP synthases on the membrane surface.

RESPONSE: Thank you for the comment. We have removed this plot to avoid confusion. The plot was originally included to demonstrate the predicted areal density of the ATP synthase enzyme was highly consistent with the experimental values. The experimental values did not include protein abundance data for growth rates higher than mu max which prevents the analysis suggested in the comment.

Fig. 4A and B - In contrast to what is said in the caption, I fail to see any inflection point in the P/O data, experimental or modeled. Furthermore, the model appears to disagree with the data as growth slows down - an increase in P/O is predicted, but the data is flat (or drops slightly).

RESPONSE: Thank you for the comment. We have replotted the analysis of the MG1655 and NCM3722 P/O data as a function of growth rate. The new analyses include the testing of alternative SA:V ratios (Figure 6E) on the P/O number as well as an experimental analysis of different empirical fits to the experimental data (supplementary data S7). The model is in fact consistent with experimental data. The P/O number inflection point quantifies where the available surface area becomes limiting and predicts when the cell will adopt an overflow metabolism which is consistent with experimental data. At slower growth rates, there is an excess of available surface area, therefore the cell does not use an overflow metabolism, and the cell can utilize some of the excess surface area to express alternative transporters in a bet hedging strategy. We have now included proteomics data for alternative ABC transporters as a function of growth rate. At low growth rates, the areal abundance of unused transporters increases, which is consistent with the predictions (Figure 3C).

Fig. 4E - Here, the authors randomly perturb the model parameters and show that the predictions do not change -- on average. Given how poorly some kinetic parameters are determined, rather than randomly perturbing parameters and taking the average, the Authors should instead study how sensitive the predictions are to changes in the value of individual parameters. For example: how are predictions affected if the ATP synthase turnover number is doubled?

Would the predictions be the same after compensating by tweaking the fMSA parameter? What about other respiratory enzymes? How would the “experimental” P/O values be affected? Such sensitivity analysis would be much more meaningful than what is presented in this panel.

RESPONSE: Thank you for the comment. We have added four additional sensitivity analyses to the document to further bolster our theory. We have perturbed single enzymes +/- 35% and quantified the effect on the network to produce maintenance energy which each of the perturbations being tested 100 times with independent perturbations in the experimental consensus fluxes. For example, decreasing the ATP synthase Kcat by 35% resulted in a 25% decrease in the capacity of the model to produce ATP. This is documented in the discussion section. We have perturbed the membrane crowding parameter (fMSA) from 0.02 to 0.2 to determine the effect on predicted NCM3722 maximum growth rate (Figure 6D), perturbed the NCM3722 SA:V ratios to determine the effect on predicted maximum growth rate (Figure 6D), perturbed the NCM3722 SA:V available to assess the effect on the predicted P/O number (Figure 6E), and we have perturbed the NCM3722 SA:V ratios to predict when the cell would be expected to transition to an overflow metabolism (Figure 6E). In all cases the theory is robust to these perturbations highlighting the importance of each of these critical parameters on predictions.

Minor:

“Strain MG1655 has an experimental mu_max of 0.69/h while the available sMSA could support a specific growth rate of 1.4/h and it has an experimental biomass yield on glucose of 0.4 g/g while it is theoretically as possible to support yields as high as 0.7 g/g”. These numbers are a bit misleading, as it is clear from Fig. 2A itself - these numbers can only be achieved when the cell is producing no energy (q_atp=0). At the very least, the energy necessary for synthesizing proteins should be taken into account, as it would restrict the solution space considerably.

RESPONSE: Thank you for the comment. The simulations do account for the energetic costs to produce amino acids and to polymerize the amino acids into proteins. The plot quantifies the influence of the assumed maintenance energy fluxes (which are independent of and in addition to the biosynthetic costs) on predicted phenotype. The current work addresses maintenance energy in a novel manner by setting constraints based on measurable SA:V ratios and protein crowding as compared to the typical FBA approach of simply assuming a value that forces the predictions and experimental data to align.

Fig. 4F, caption is mislabeled

RESPONSE: Thank you for the careful reading of the document. We have corrected the numbering on the figure caption.

## List of abbreviations

SA:V: surface area to volume
*f_MSA_*: fraction of membrane surface area
*MRE_i_*: membrane real estate for enzyme *i*
*sMSA*: specific membrane surface area (*nm*^2^ (*gcdw*)^-1^)
*sMSAc-FBA*: specific membrane surface area constrained flux balance analysis
*q_ATP_*: specific ATP production rate (*mmol ATP* (*gcdw h*)^-1^)
*q_glucose_*: specific glucose consumption rate (*mmol glucose* (*gcdw h*)^-1^)
*q_acetate_*: specific acetate secretion rate (*mmol acetate* (*gcdw h*)^-1^)
P/O: phosphate (ATP bonds) to oxygen (2 electrons) ratio
ETC: electron transport chain
μ_max_: maximum specific growth rate
Nuo: NADH dehydrogenase enzyme complex I
Ndh II NADH: dehydrogenase enzyme complex II
Cyo: cytochrome *bo*
Cyd: cytochrome *bd*
PtsG: phosphotransferase system glucose
GAM: growth associated maintenance energy
NGAM: nongrowth associated maintenance energy
*k_cat_*: enzyme catalytic parameter

